# Dynamic *Runx1* chromatin boundaries affect gene expression in hematopoietic development

**DOI:** 10.1101/2021.05.14.444178

**Authors:** Dominic D.G. Owens, Giorgio Anselmi, A. Marieke Oudelaar, Damien J. Downes, Alessandro Cavallo, Joe R. Harman, Ron Schwessinger, Akin Bucakci, Lucas Greder, Sara de Ornellas, Danuta Jeziorska, Jelena Telenius, Jim R. Hughes, Marella F.T.R. de Bruijn

## Abstract

The transcription factor RUNX1 is a critical regulator of developmental hematopoiesis and is frequently disrupted in leukemia. *Runx1* is a large, complex gene that is expressed from two alternative promoters under the spatiotemporal control of multiple hematopoietic enhancers. To dissect the dynamic regulation of *Runx1* in hematopoietic development, we analyzed its three-dimensional chromatin conformation in mouse embryonic stem cell (ESC) differentiation cultures. *Runx1* resides in a 1.1 Mb topologically associating domain (TAD) demarcated by convergent CTCF motifs. As ESCs differentiate to mesoderm, chromatin accessibility, *Runx1* enhancer-promoter (E-P) interactions, and CTCF-CTCF interactions increased in the TAD, along with initiation of *Runx1* expression from the P2 promoter. Differentiation to hematopoietic progenitor cells was associated with the formation of tissue-specific sub-TADs over *Runx1*, a shift in E-P interactions, P1 promoter demethylation, and robust expression from both *Runx1* promoters. Deletions of promoter-proximal CTCF sites at the sub-TAD boundaries had no obvious effects on E-P interactions but led to partial loss of domain structure, mildly affected gene expression, and delayed hematopoietic development. Together, our analyses of gene regulation at a large multi-promoter developmental gene revealed that dynamic sub-TAD chromatin boundaries play a role in establishing TAD structure and coordinated gene expression.

## Introduction

Runx1/AML1 is a member of the RUNX family of transcription factors (TFs), which are key to many developmental processes (de Bruijn and Dzierzak 2017, Gao et al. 2018, Levanon and Groner 2004). *Runx1* is best known for its critical role in the *de novo* generation of the hematopoietic system and maintenance of normal hematopoietic homeostasis (de Bruijn and Dzierzak 2017, Levanon and Groner 2004, Yzaguirre et al. 2017). Disruption of *RUNX1* in humans leads to several hematopoietic disorders, including acute myeloid leukemia (Sood et al. 2017) and familial platelet disorder with associated myeloid malignancy (FPD-AML)(Bellissimo and Speck 2017). All members of the RUNX family bind the same canonical DNA motif (YGYGGT) and their tissue-specific functions are thought to be governed largely by their specific expression patterns (Levanon et al. 2001). *Runx1* transcription is tightly regulated, with changes in gene dosage and expression level affecting both the spatiotemporal onset of hematopoiesis and hematopoietic homeostasis (Cai et al. 2000, Lie et al. 2018, Song et al. 1999, Wang Q. et al. 1996). *Runx1* is transcribed from two alternative promoters, the distal P1 and proximal P2 that are differentially regulated and generate different transcripts and protein products (Ghozi et al. 1996, Telfer and Rothenberg 2001, and reviewed in de Bruijn and Dzierzak (2017)). During hematopoietic development, *Runx1* expression initiates from the P2 promoter and gradually switches to the P1 promoter with the majority of adult hematopoietic cells expressing P1-derived *Runx1* (Bee et al. 2009b, Bee et al. 2010, North et al. 1999, Sroczynska et al. 2009a, Telfer and Rothenberg 2001). The *Runx1* promoters do not confer tissue specificity and several distal *Runx1* cis-elements have been identified that mediate reporter gene expression in transient transgenic embryos in *Runx1*-specific spatiotemporal patterns (Bee et al. 2010, Marsman et al. 2017, Ng et al. 2010, Nottingham et al. 2007, Schutte et al. 2016). However, the combinatorial regulation of *Runx1* at different stages of hematopoiesis is currently unclear, as are the mechanisms through which the tight and dynamic spatiotemporal control of *Runx1* expression is achieved. A better understanding of *Runx1* regulation may yield insights into potential avenues for therapeutic targeting of *RUNX1* in a variety of hematological disorders, as recently highlighted by growth inhibition in a leukemia cell line upon loss of the *Runx1* +23 enhancer (Mill et al. 2019).

The 3D conformation of DNA in structures such as topologically associating domains (TADs) delimit the activities of enhancers *in vivo* (Hanssen et al. 2017, Lettice et al. 2011, Lupianez et al. 2015, Symmons et al. 2014). Specific interactions achieved through chromatin folding, particularly enhancer-promoter (E-P) interactions, are thought to be a key component of spatiotemporal gene regulation (Oudelaar and Higgs 2021, Schoenfelder and Fraser 2019). Many insights into principles of transcriptional regulation have come from studying a few relatively small gene loci, including the globin genes. However, genes encoding developmentally important TFs, including LIM-Homeobox, Hox, Eomes, Sox, and Sonic Hedgehog (Shh), often lie in larger regulatory domains and are frequently flanked by gene deserts (Ovcharenko et al. 2005). Indeed, studies of developmental regulators, such as Shh, have revealed exceptionally long-range enhancer-promoter interactions (Lettice et al. 2002), suggesting that specific regulatory mechanisms may be at play at these large developmental loci in addition to the basic regulatory principles established at smaller genes. One aspect that remains unclear is whether large-scale chromatin conformation changes may be required to coordinate complex developmental expression patterns at larger genes. A known factor important for the regulation of chromatin conformation is CCCTC-Binding factor (CTCF) (Braccioli and de Wit 2019). CTCF, along with the loop extruding factor cohesin, mediates the establishment and maintenance of both E-P interactions and TAD structure (Nora et al. 2017, Rao et al. 2017, Schwarzer et al. 2017). Interestingly, *Runx1* was mis-regulated in zebrafish after perturbation of CTCF/cohesin (Horsfield et al. 2007, Marsman et al. 2014, Mazzola et al. 2020), suggesting that *Runx1* regulation may depend on chromatin structure. Elucidating *Runx1* transcriptional regulatory mechanisms is expected to contribute to a better understanding of the chromatin conformation changes employed by complex multi-promoter genes during development.

Here, we characterized the *Runx1* chromatin landscape in four dimensions, i.e. in 3D space over time, in an *in vitro* mouse ESC differentiation model of developmental hematopoiesis. Using high-resolution chromosome conformation capture (Tiled-C) (Oudelaar et al. 2020), we report the presence of a pre-formed 1.1 Mb TAD spanning the *Runx1* locus in mouse ESCs that is conserved in human and forms prior to gene activation. Upon differentiation, accessible chromatin sites emerged within the TAD over known enhancers, CTCF sites, and novel candidate cis-regulatory elements. These regions interacted with the *Runx1* promoters in a developmental stage-specific manner. Notably, an increased interaction of the P1 and P2 promoters within cell type-specific *Runx1* sub-TADs was seen. These sub-TADs were bounded by highly conserved promoter-proximal CTCF sites, the role of which is poorly understood. Here, we used a machine learning approach and CRISPR/Cas9-mediated deletion to examine the importance of promoter-proximal CTCF binding for the *Runx1* chromatin landscape. Deletion of either the *Runx1* P1 or P2 promoter-proximal CTCF site partially disrupted TAD structure, while E-P interactions appeared unaffected. *Runx1* levels showed a decreased trend at the mesoderm stage, concomitant with significant changes in mesodermal gene expression indicative of a delay in hematopoietic differentiation. Together, we find that sub-TAD chromatin boundaries form dynamically within the large and complex *Runx1* regulatory domain during differentiation and are involved in coordinating gene expression and hematopoietic differentiation.

## Results

### *Runx1* lies in a conserved TAD which forms prior to gene activation

To investigate dynamic changes in 3D chromatin confirmation in the *Runx1* locus (schematically represented in Figure 1A) during hematopoietic development, we used the in vitro mouse ESC (mESC) differentiation model that recapitulates the *de novo* generation of hematopoietic progenitor cells (HPCs) from mesoderm as it occurs in the embryo, including the endothelial-to-hematopoietic transition (EHT) specific to development (adapted from Sroczynska et al. (2009b)). In this model, we assessed chromatin conformation (Tiled-C), along with gene expression (poly(A)-minus RNA-seq to capture nascent transcripts) and chromatin accessibility (ATAC-seq) over hematopoietic differentiation (Figure 1B). Flk1^+^ mesodermal cells were isolated by flow cytometry from day 4 embryoid body (EB) cultures. Upon further differentiation in EHT media these gave rise to phenotypic HPCs with blood progenitor morphology and *in vitro* clonogenic potential (Figure 1B, C, Supp Figure 1). Gene expression analysis of mESCs, Flk1+ mesoderm and emerging CD41^+^ CD45^-^ Runx1^+^ HPCs reflected the developmental trajectory as visualized in a Principal Component Analysis (PCA) plot (Figure 1D). This was accompanied by silencing of pluripotency genes (*Pof5f1, Sox2, Nanog*), transient expression of mesodermal genes (*Kdr, Eomes, T*), and increasing levels of hematopoiesis-associated genes (*Pecam1, Tal1, Gfi1b, Meis1, Itga2b*) (Figure 1E). *Runx1* expression was initiated in Flk1+ mesoderm and increased in HPCs (Figure 1E; Sroczynska et al. (2009a)).

**Figure 1.**
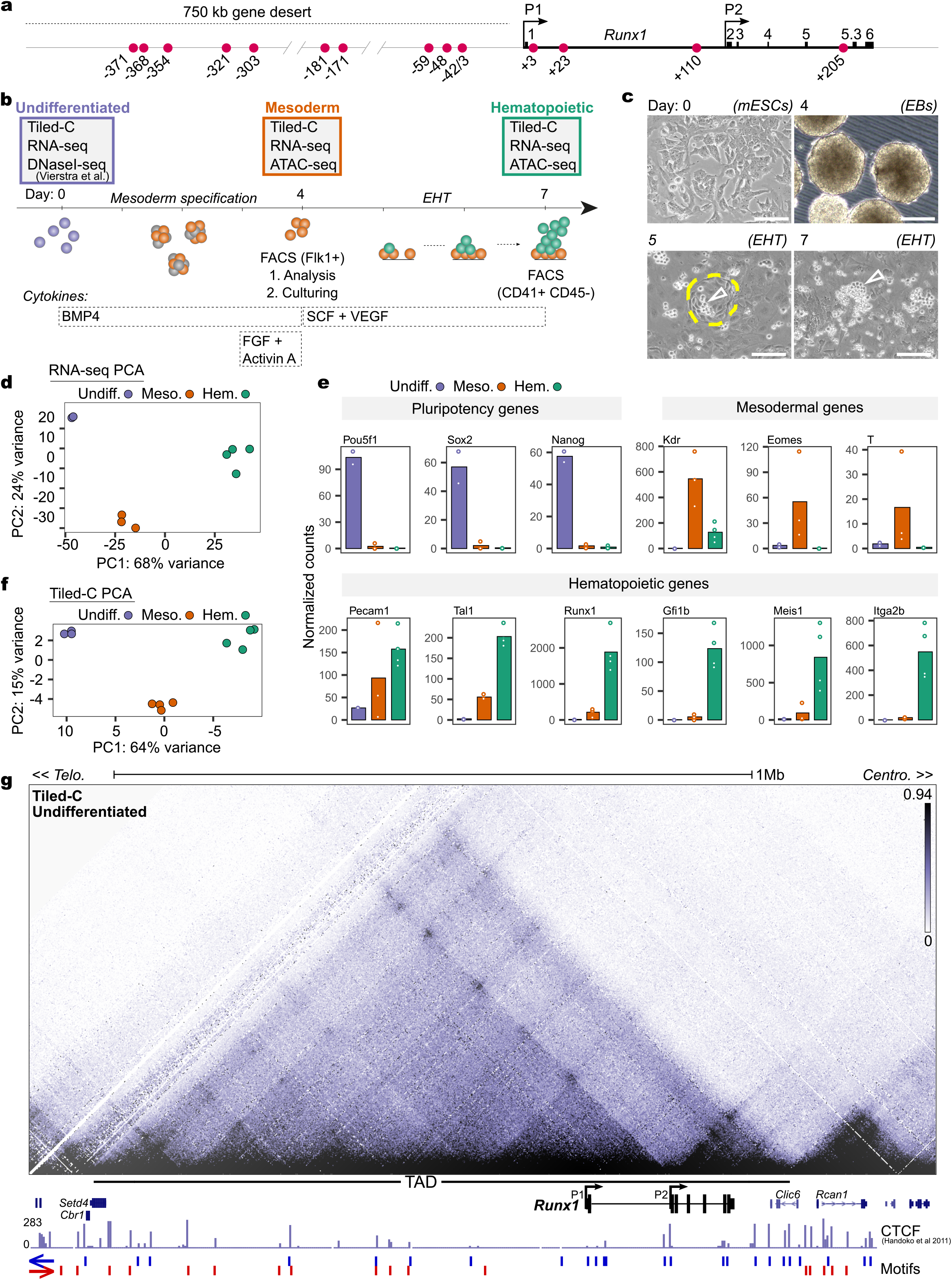
*Runx1* resides within a topologically associating domain (TAD) in undifferentiated cells. **A)** Schematic of the *Runx1* locus on mouse chromosome 16, with *Runx1* proximal (P2) and distal (P1) promoters, exons, and adjacent gene desert labelled. Previously identified enhancers are indicated by red circles that are numbered according to the distance (in kb) from the *Runx1* start codon in exon 1 (Bee et al. 2009a, Bee et al. 2009b, Bee et al. 2010, Ghozi et al. 1996, Marsman et al. 2017, Miyoshi et al. 1995, Ng et al. 2010, Nottingham et al. 2007, Schutte et al. 2016). **B)** Schematic of seven-day differentiation protocol with cytokines and markers used for isolation of cells by FACS indicated. EHT = endothelial-to-hematopoietic transition. DNaseI-seq data in mESC was previously published (Vierstra et al. 2014). **C)** Bright-field images of different stages of *in vitro* differentiation. Colonies of hemogenic endothelial (HE) cells are outlined with dashed yellow lines and clusters of emerging hematopoietic progenitors are indicated by hollow white arrowheads. Scale bars = 200 *µ*m. **D)** Principal component analysis (PCA) of individual poly(A) minus RNA-seq replicates colored by cell type. **E)** Plot of normalized counts of lineage marker gene expression across differentiation. Undifferentiated n=2, mesoderm n=3, hematopoietic n=4. **F)** PCA of individual Tiled-C replicates colored by cell type. **G)** Tiled-C matrix at 2 kb resolution for undifferentiated mESCs. Matrix is a merge of three independent replicates (n=3). Interactions are visualized with a threshold at the 94^th^ percentile. *Runx1* promoters (P1 and P2), neighboring genes, the adjacent gene desert, and approximate location of the 1.1 Mb *Runx1* TAD are labelled. Publicly available CTCF ChIP-seq in E14 mESCs (Handoko et al. 2011) was reanalyzed and the orientation of CTCF motifs identified *de novo* under CTCF peaks is indicated.

To generate high resolution chromatin conformation maps of the *Runx1* regulatory domain, we performed Tiled-C, a targeted method that generates Hi-C-like data at specific loci (Oudelaar et al. 2020), with probes against all *DpnII* fragments in a 2.5 Mb region centered on *Runx1*. PCA of individual Tiled-C replicates showed a clear developmental trajectory (Figure 1F, Supp. Figure 2) similar to that seen based on gene expression analysis (cf. Figure 1D), demonstrating that *Runx1* exhibits reproducible dynamic chromatin conformation changes during differentiation. *Runx1* resides within a 1.1Mb TAD in mESCs (Figure 1G, mm9 chr16:92496000-93617999) that extends to encompass the upstream 750 kb gene desert, with the *Setd4* and *Cbr1* genes at its telomeric end, and *Clic6* at its centromeric end (Figure 1G). Using previously published CTCF occupancy data from mESCs (Handoko et al. 2011), thirty-one binding sites were identified within the *Runx1* TAD (Figure 1G). MEME analysis (Bailey and Elkan 1994) identified core CTCF binding motif location and orientation underlying CTCF peaks and revealed a predominant convergence of CTCF motifs—with primarily centromeric oriented motifs near the *Setd4* telomeric end of the *Runx1* TAD, and telomeric oriented motifs primarily at the *Clic6* centromeric end (Figure 1G). Together, this shows that the *Runx1* regulatory domain is established prior to gene expression, likely in a CTCF-dependent manner.

### Mesodermal differentiation is accompanied by increased interactions between *Runx1* P2 and enhancers in the gene desert and *Runx1* gene body

Upon transition from mESC to Flk1^+^ mesoderm we observed increased chromatin accessibility in the *Runx1* TAD and low levels of transcription primarily from the *Runx1* P2 promoter (Figure 2A, B). At this mesodermal stage, we identified thirty-three open chromatin sites that are absent from mESCs (DNaseI-seq; Vierstra et al. (2014)) (Figure 2A, Supp. Table 1, adjusted p-value < 0.05). Ten peaks corresponded to previously identified enhancers (−327, -322, -303, -181, -171, -59, +3, +23, +48, +110) (Bee et al. 2010, Cauchy et al. 2015, Cheng et al. 2018, Fitch et al. 2020, Harland et al. 2021, Marsman et al. 2017, Ng et al. 2010, Nottingham et al. 2007, Ortt et al. 2008, Schutte et al. 2016) (Supp. Table 2), while twenty-four peaks unique to mesoderm did not overlap with any known regulatory elements (Supp. Table 1). Interactions between CTCF sites also increased at the mesodermal stage, particularly between the two boundaries of the *Runx1* TAD (Figure 2C, D, Kruskal-Wallis and Dunn’s test, adjusted p = 0.003), while insulation of the main TAD (a measure of intra-TAD interactions) decreased slightly (Figure 2E, Kruskal-Wallis and Dunn’s test, adjusted p = 1.4×10^−135^). To determine promoter-specific enhancer interactions, we compared virtual Capture-C profiles across the *Runx1* locus using the P2 and P1 promoters as viewpoints (Figure 2F). We observed an overall increase of E-P2 interactions in mesoderm compared to mESCs (Figure 2F, Kruskal-Wallis and Dunn’s test, adjusted p = 0.005), specific increased interactions between the P2 promoter and the -327, -322, -303, -181, -171 enhancer elements in the gene desert, and the +3, +23 and +110 enhancers in the *Runx1* gene body (Figure 2G). In contrast to the P2, the P1 promoter did not show a significant overall increase in interactions with enhancers in mesodermal cells (Figure 2F, Kruskal-Wallis and Dunn’s test, adjusted p = 0.9), in line with the absence of P1-derived *Runx1* expression (Figure 2B). However, a slight specific increase was seen in interactions between P1 and elements -181 and -171 in the gene desert, and elements +48, +110 within *Runx1* intron 1, and with the P2 could be seen (Figure 2G). Together, these data indicate that early spatiotemporal control of *Runx1* expression at the onset of hematopoiesis is associated with increased CTCF interactions, reduced TAD insulation, and may be mediated by regulation of interactions between specific enhancer elements and both the P2 and P1 promoters.

**Figure 2.**
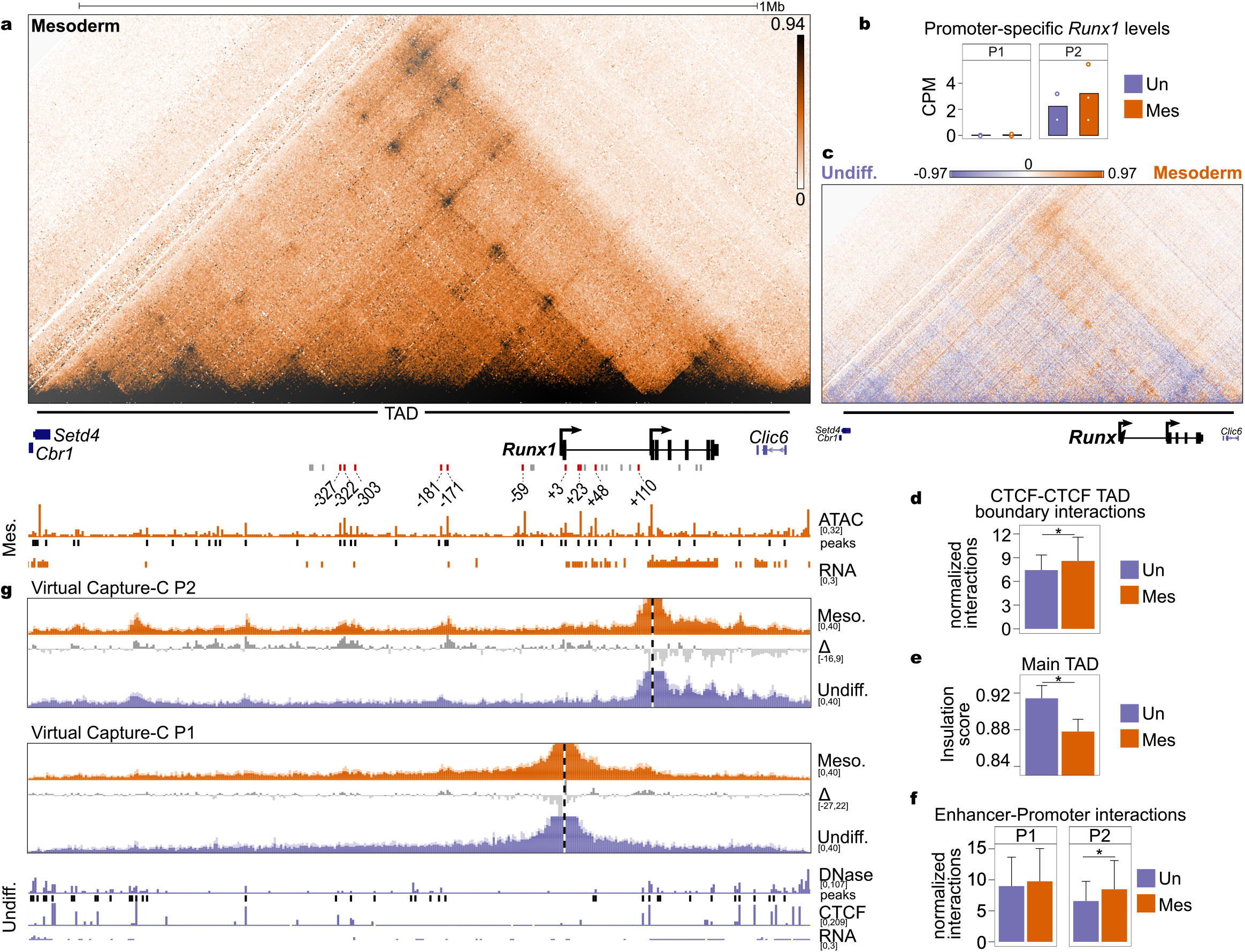
Early hematopoietic differentiation leads to increased enhancer*-Runx1* P2 interactions. **A)** Tiled-C matrix from mesoderm (2 kb resolution, threshold at the 94^th^ percentile, n=4). *Runx1* promoters and location of *Runx1* TAD are labelled below the matrix. RPKM-normalized ATAC-seq (n=4 replicates from two independent experiments) track is shown with called peaks (adjusted p < 0.05). CPM-normalized poly(A)-minus RNA-seq (n=3) is shown. Previously published enhancer regions are indicated. Enhancer regions that are accessible in mesoderm are shown as red bars and numbered according to their distance from the *Runx1* start codon in exon 1. Enhancers that did not overlap ATAC-seq peaks are identified by grey bars. **B)** Promoter-specific *Runx1* levels in undifferentiated and mesoderm cells. **C)** Subtraction of normalized Tiled-C matrices between undifferentiated and mesoderm. The matrix is a subtraction of the signal between two merged matrices (undifferentiated n=3, mesoderm n=4, 2 kb resolution, threshold at +97^th^ and -97^th^ percentile). **D)** Quantification of interactions between the four outermost CTCF peaks at the edges of the TAD (*, Kruskal-Wallis and Dunn’s test, adjusted p = 0.003). Each bar represents the median value and error bars show the interquartile range. **E)** Insulation score (intra-TAD interaction ratio) of the main *Runx1* TAD (*, Kruskal-Wallis and Dunn’s test, adjusted p = 1.4×10^−135^). Each bar represents the median insulation score across bins and error bars show the interquartile range. **F)** Quantification of total interactions from the viewpoint of each promoter with all previously published enhancers (Supp. Table 2). Each bar represents the median contacts and error bars show the interquartile range. **G)** Virtual Capture-C profiles (obtained from Tiled-C data, see methods) from the viewpoint of both *Runx1* promoters in undifferentiated mESCs (blue tracks) and mesodermal cells (orange tracks). *Runx1* promoters (P1 and P2) are indicated by a vertical dashed line. Dark colors represent the mean reporter counts in 2 kb bins (undifferentiated n=3, mesoderm n=4) normalized to the total *cis*-interactions in each sample. Standard deviation is shown in the lighter color. Subtractions between two cell types as indicated are shown as grey tracks. DNaseI-seq (Vierstra et al. 2014), DNaseI peaks, CTCF ChIP-seq (Handoko et al. 2011), and RNA-seq from undifferentiated mESCs are indicated in blue below Capture-C tracks.

### Increased *Runx1* expression upon hematopoietic differentiation is associated with P1 activation, a shift in E-P interactions and sub-TAD reinforcement

Differentiation of hematopoietic-fated mesoderm into HPCs is accompanied by increased expression from both *Runx1* promoters, with a three-fold higher expression from P2 than P1 (Figure 3A, B). Compared to mesoderm, HPCs show dynamic shifts in chromatin accessibility in the *Runx1* TAD. ATAC-seq peaks were gained in the gene body at the +204 and -42 enhancers, lost at +48, -171, -181, -303, -322, -328 and other sites in the gene desert, and maintained at the +3, +23, +110 enhancers in HPCs (Figure 3A, MACS2 adjusted p < 0.05, Supp. Table 1). Insulation of the main TAD slightly decreased further as cells differentiated from mesoderm to HPCs (Figure 3C, Kruskal-Wallis and Dunn’s test, adjusted p = 2.1×10^−7^), while CTCF-CTCF interactions between the boundaries of the TAD were not different from mesoderm (Figure 3C, Kruskal-Wallis and Dunn’s test, adjusted p = 0.13). Alongside these changes at the main TAD-level, two sub-TADs spanning the *Runx1* gene were strengthened significantly in HPCs (Figure 3D, E, Kruskal-Wallis and Dunn’s test, P1-P2 sub-TAD p = 3.1×10^−43^ and P2-3’ sub-TAD p = 6.6×10^−13^). We next compared how specific E-P interactions changed between mesoderm and HPCs using virtual Capture-C plots of the Tiled-C data. Compared to mesoderm, total E-P interactions increased in HPCs for the P2 promoter and there was a trend of increased overall E-P interactions for P1 (Figure 3F, Kruskal-Wallis and Dunn’s test, P2: adjusted p = 0.005, P1: adjusted p = 0.06). Both P1 and P2 showed specific increases in E-P interactions with -59, -43, -42, +23, +110 enhancers (Figure 3G), while interactions with elements extending further in the gene desert were lost. Therefore, differentiation to HPCs is associated with specific E-P interactions primarily within HPC-specific sub-TADs that span the *Runx1* gene.

**Figure 3.**
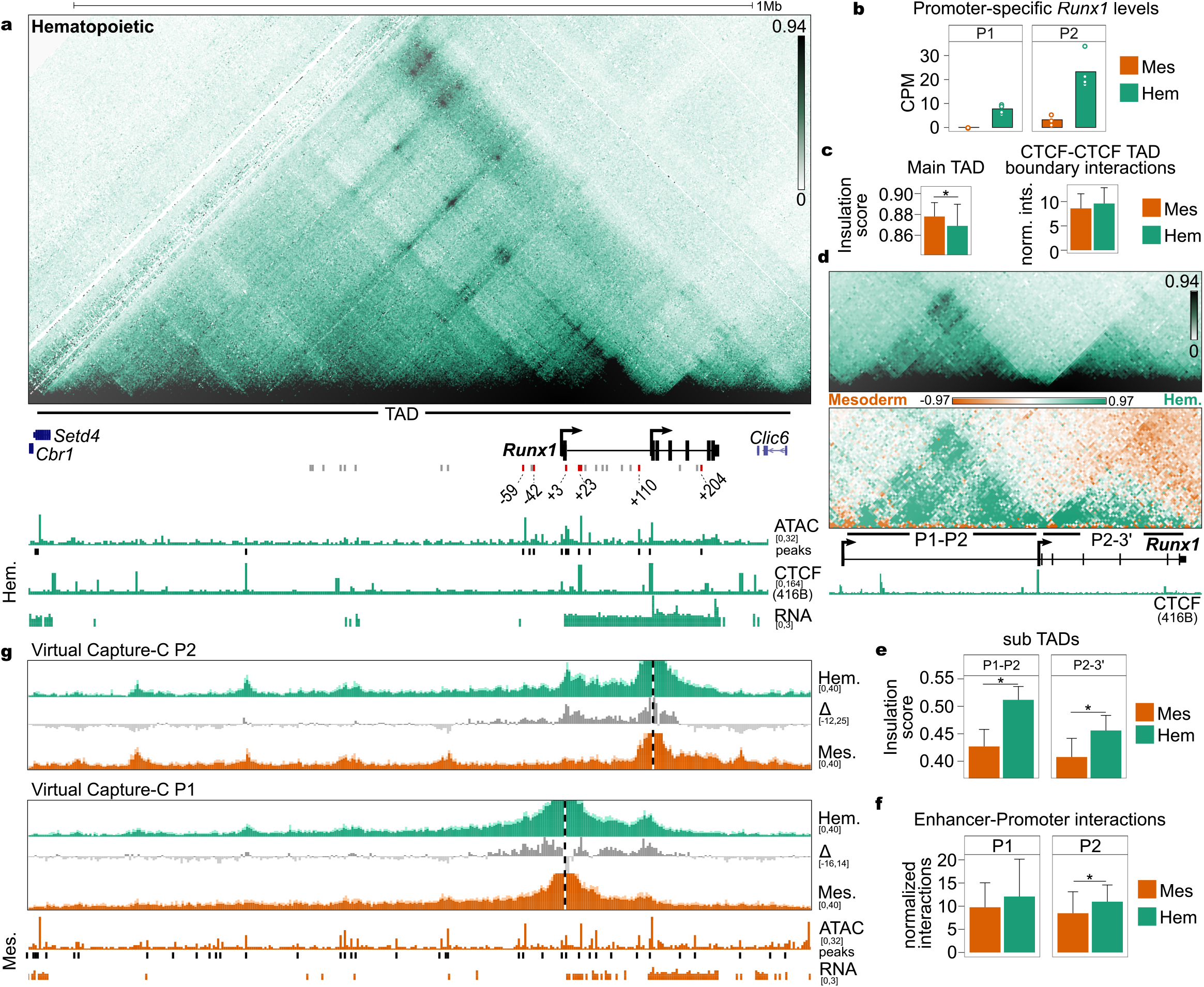
EHT progression is associated with sub-TAD reinforcement, increased *Runx1* expression and P1 activation. **A)** Tiled-C matrix from HPCs (2 kb resolution, threshold at the 94^th^ percentile, n=4). *Runx1* promoters and location of *Runx1* TAD are labelled below the matrix. RPKM-normalized ATAC-seq (n=2 replicates from one experiment) track is shown with called peaks (adjusted p < 0.05). CPM-normalized poly(A)-minus RNA-seq (n=3) is shown. Previously published enhancer regions are indicated. Enhancer regions that are accessible in HPCs are shown as red bars and numbered according to their distance from the *Runx1* start codon in exon 1. Enhancers that did not overlap ATAC-seq peaks are identified by grey bars. **B)** Promoter-specific *Runx1* levels in mesoderm and HPCs. **C)** Left, insulation score (intra-TAD interaction ratio) of the main *Runx1* TAD (*, Kruskal-Wallis and Dunn’s test, adjusted p = 2.1×10^−7^). Right, quantification of interactions between the four outermost CTCF peaks at the edges of the TAD. Each bar represents the median value and error bars show the interquartile range. **D)** Top, zoom of Tiled-C data at 2 kb resolution with a threshold at 94^th^ percentile. Below, subtraction of normalized Tiled-C matrices between mesoderm and HPCs. The matrix is a subtraction of the signal between two merged matrices (n=4, 2 kb resolution, threshold at +97^th^ and -97^th^ percentile). **E)** Insulation scores (intra-TAD interaction ratio) of the two *Runx1* sub-TADs (*, Kruskal-Wallis and Dunn’s test, P1-P2 TAD p = 3.1×10^−43^ and P2-3’ TAD p = 6.6×10^−13^). Each bar represents the median insulation score across bins and error bars show the interquartile range. **F)** Quantification of total interactions from the viewpoint of each promoter with all previously published enhancers (Supp. Table 2). Each bar represents the median of contacts and error bars show the interquartile range. **G)** Virtual Capture-C profiles (obtained from Tiled-C data, see methods) from the viewpoint of both *Runx1* promoters in mesoderm (orange tracks) and HPCs (green tracks). *Runx1* promoters (P1 and P2) are indicated by a vertical dashed line. Dark colors represent the mean reporter counts in 2 kb bins (n=4) normalized to the total *cis*-interactions in each sample. Standard deviation is shown in the lighter color. Subtractions of the signal between two cell types as indicated are shown as grey tracks. ATAC-seq and peaks and RNA-seq from mesoderm are indicated in orange below Capture-C tracks.

### *Runx1* promoter-proximal CTCF sites play a role in establishing *Runx1* chromatin architecture but not E-P interactions

The sub-TAD spanning the first intron of *Runx1* had boundaries that correspond to the P1 and P2 promoters. As both promoters have telomeric orientated CTCF sites less than 2kb upstream of the transcription start site (TSS; Figure 3A, D), and CTCF is associated with sub-TAD and TAD boundaries (Braccioli and de Wit 2019, Dixon et al. 2012), we performed CTCF ChIP-seq in the 416B HPC cell line to determine if the observed changes in sub-TAD structure may be associated with differential CTCF binding in HPCs versus mESCs. Like mESC-derived HPCs, 416B cells HPCs express *Runx1* from both the P1 and P2 promoter (Figure 3A, Supp. Figure 3A). Interestingly, while the majority of CTCF sites in the *Runx1* TAD were bound at similar levels in HPCs and mESCs, an increase in CTCF binding was seen in HPCs at the P1-proximal CTCF site, and at the +23 enhancer (Supp. Figure 3A). We next examined what the mechanism underlying the differential CTCF binding could be. DNA CpG dinucleotide methylation has been suggested to modulate dynamic CTCF binding (Bell and Felsenfeld 2000, Canzio and Maniatis 2019, Flavahan et al. 2016, Hashimoto et al. 2017, Schuijers et al. 2018, Shukla et al. 2011, Wang H. et al. 2012, Xu and Corces 2018), and is also known to be associated with promoter silencing (Curradi et al. 2002). To investigate whether P1 promoter methylation could underlie differential activation and CTCF binding during hematopoietic differentiation, we performed targeted bisulfite sequencing of *Runx1* promoters in undifferentiated mESCs and 416B HPCs. While the *Runx1* P2 promoter harbors a 2.0 kb CpG island that was hypomethylated in both cell types (Supp. Figure 3B), in contrast, *Runx1* P1 was near-completely methylated in mESCs, and became demethylated in hematopoietic cells (Supp. Figure 3B). Together, this shows that *Runx1* sub-TAD strengthening over hematopoietic differentiation is associated with *Runx1* P1 promoter demethylation and increased CTCF binding at promoter-proximal sites.

Genome-wide, we observed a significant enrichment of CTCF binding close to active promoters in both mESCs and HPCs (< 5kb from TSS; Supp. Figure 3C, D, Chi-square test, p < 1×10^−10^). The role of promoter-proximal CTCF has not been widely explored. To examine whether the deeply conserved P1 and/or P2 promoter proximal CTCF sites (Figure 4A), hereon referred to as P1-CTCF and P2-CTCF, may play a role in establishing the dynamic *Runx1* sub-TADs in HPCs we first utilized a novel deep learning approach (deepC; Schwessinger et al. (2020)) to predict chromatin interactions at the *Runx1* locus in mESCs. The deepC model was trained on Hi-C data from mESCs (Bonev et al. 2017), withholding mouse chromosome 16 containing *Runx1*. The overall *Runx1* TAD predicted by deepC agreed well with the TAD observed in Tiled-C data from mESCs (Supp. Figure 4A). In silico deletion of the CTCF site proximal to the P2 promoter was predicted to reduce the stripe of interactions emanating from this site into the gene desert, and to increase interactions across the boundary in mESCs (Supp. Figure 4B). In contrast, deletion of the P1 proximal CTCF was predicted have little effect on chromatin interactions in mESCs (Supp. Figure 4C).

**Figure 4.**
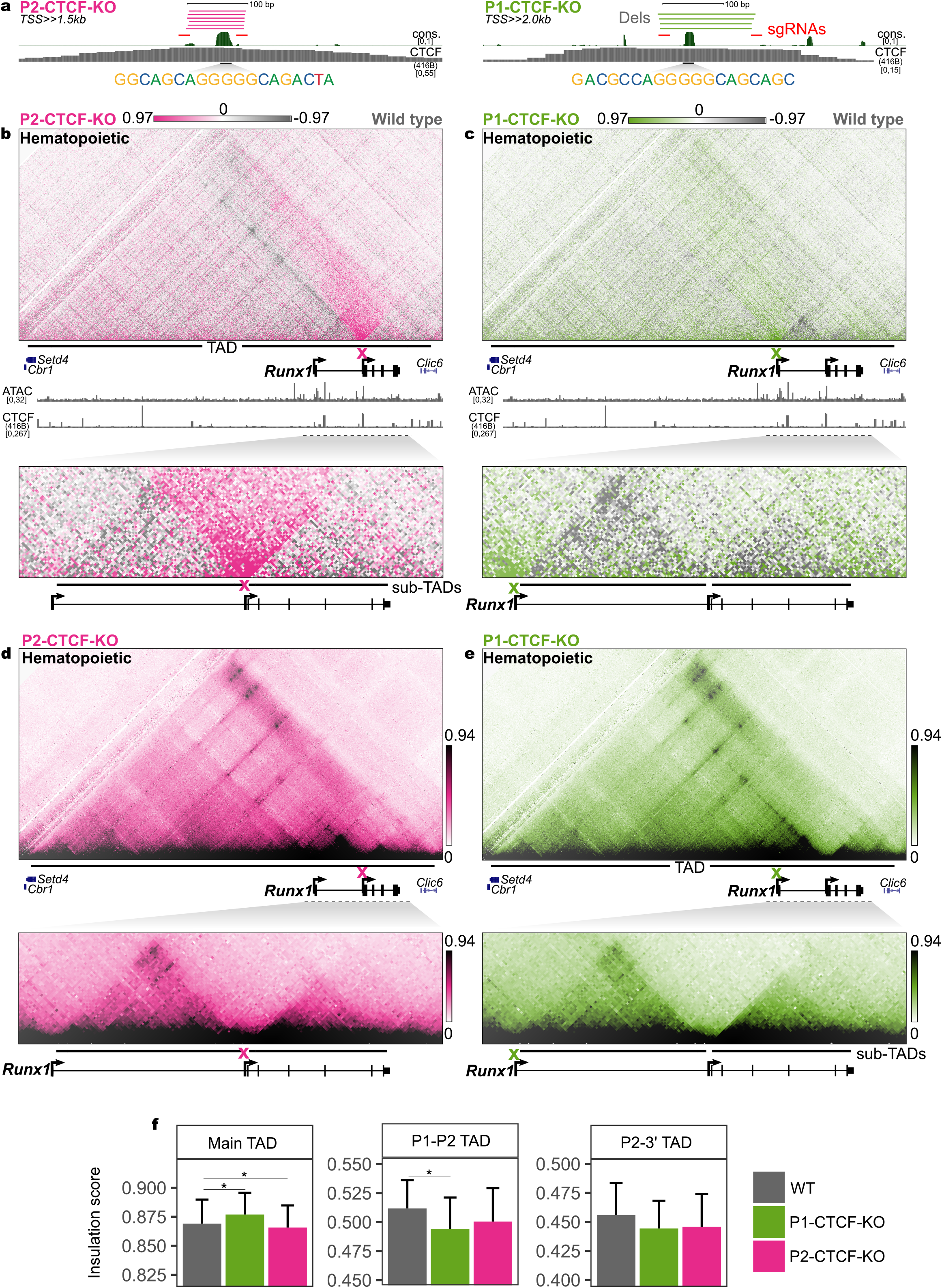
*Runx1* promoter-proximal CTCF sites play a role in establishing *Runx1* chromatin architecture. **A)** Schematic of *Runx1* P1 and P2 promoter-proximal CTCF sites and CRISPR/Cas9 strategies to delete them. Distance to *Runx1* transcription start sites is indicated. Vertebrate conservation (phastCons), CTCF occupancy in 416B HPCs, core motif sequence, single guide (sg)RNA, and deletion alleles (dels) are indicated. **B, C)** Subtraction of Tiled-C matrices between P2-CTCF-KO (B) P1-CTCF-KO (C) and wild type hematopoietic cells is shown (2 kb resolution, threshold at +/-97^th^ percentile, n=4). Locations of CTCF site deletions are indicated by a pink and green cross. RPKM-normalized ATAC-seq in wild type HPCs (n=2 replicates from one experiment) and CTCF occupancy in 416B cells is shown. The locations of the main *Runx1* TAD and sub-TADs are indicated. **D, E)** Tiled-C matrix from P2-CTCF-KO (D) and P2-CTCF-KO (E) (2 kb resolution, threshold at 94^th^ percentile, n=4). **F)** Insulation scores (intra-TAD interaction ratio) for main *Runx1* TAD and sub-TADs in wild type, P1-CTCF-KO, and P2-CTCF-KO HPCs (*, Kruskal-Wallis and Dunn’s test, p < 0.05). Each bar represents the median insulation scores across bins and error bars show the interquartile range.

Next, we determined the impact of promoter-proximal CTCF site deletion on chromatin conformation experimentally by Tiled-C. We generated P1-CTCF-KO and P2-CTCF-KO mESC clones using CRISPR-Cas9; these lacked the entire CTCF site, including the core motif, but retained nearby conserved sequences (Figure 4A, Supp. Figures 5-7). Hematopoietic differentiation in the KO clones was unaffected (three independent mESCs clones analyzed each for the P1-CTCF-KO and P2-CTCF-KO; Supp. Figure 8) and Tiled-C was performed on undifferentiated mESCs, Flk1+ mesoderm, and HPCs. PCA showed that P1-CTCF-KO and P2-CTCF-KO cells clustered along the same developmental trajectory as wild type mESC differentiation cultures (Supp. Figures 9 and 10). Strikingly, CTCF-CTCF interactions were reduced across the entire TAD in P2-CTCF-KO HPCs, in agreement with the deepC prediction and consistent with P2-CTCF forming an insulated boundary (Figure 4B, upper panel). Tiled-C in P1-CTCF-KO mESCs also agreed with the deepC predictions, with loss of the P1-CTCF site showing no effect compared to wild type mESCs (Supp. Figure 4C). P1-CTCF-KO HPCs, however, exhibited a subtle decrease in interactions with CTCF sites in the gene desert (Figure 4C, upper panel), highlighting the tissue-specific nature of this CTCF site. Deletion of either P1-CTCF or P2-CTCF increased interaction frequencies between regions upstream and downstream of these sites, indicating that both CTCF sites act as boundaries (Figure 4B and C, lower panels). The main TAD and tissue-specific sub-TADs were still present in P1-CTCF-KO or P2-CTCF-KO HPCs (Figure 4D, E), though insulation scores were affected, indicating that sub-TAD boundary strengths were reduced (Figure 4F). In HPCs, both P1 and P2 promoters primarily interacted with enhancers lying within the tissue-specific sub-TADs (cf. Figure 3G). As these sub-TADs were altered upon deletion of promoter-proximal CTCF motifs, E-P interactions in P1- and P2-CTCF-KO HPCs were compared to wild type cells. Surprisingly, despite generally weaker sub-TAD interactions in P1- and P2-CTCF-KO HPCs (Figure 4A, B, and E), total E-P interactions were not significantly different compared to wild type for either promoter at any stage of differentiation (Supp. Figure 11A, B, Kruskal-Wallis test, adjusted p > 0.4) nor was any individual E-P interaction (Supp. Figure 12, Kruskal-Wallis test, adjusted p = 1.0). Together, our results show that despite perturbed chromatin architecture resulting from the absence of conserved promoter-proximal CTCF sites, specific *Runx1* E-P interactions are maintained.

### *Runx1* P2 promoter-proximal CTCF site coordinates spatiotemporal gene expression and differentiation

Since CTCF binding close to promoters genome-wide, including at *Runx1* P1, is associated with promoter activity (Supp. Figure 3C, D), and since loss of promoter-proximal CTCF sites disrupted *Runx1* chromatin architecture, the effect of promoter-proximal CTCF loss on *Runx1* expression was examined during hematopoietic differentiation of KO mESC clones. We observed a non-significant trend for reduced total *Runx1* expression in P2-CTCF-KO mesoderm compared to both wild-type and P1-CTCF-KO (Figure 5A, zoomed in graph with dashed outline, DESeq2 adjusted p = 0.6). No changes were observed in alternative P1 or P2 promoter usage after deletion of promoter-proximal CTCF sites (Figure 5B). PCA of global RNA-seq profiles across all stages and genotypes showed clustering based on cell type rather than genotype (Figure 5C). However, when considering mesoderm samples alone, P2-CTCF-KO samples formed a separate cluster (Figure 5D). Indeed, differential expression analysis revealed that, globally, 168 genes were differentially expressed between P2-CTCF-KO and wild type mesoderm (Figure 5E, DESeq2 adjusted p < 0.05, fold change > 1). Notably, expression of several mesodermal markers (including *T* and *Eomes*) was higher in P2-CTCF-KO mesoderm compared to wild type, while several hematopoietic markers were downregulated (Figure 5F, adjusted p < 0.05). GO analysis of genes downregulated by P2-CTCF-KO were associated with biological processes including “response to growth factor” and “blood vessel remodeling”, while upregulated genes were associated with terms including “mesoderm development” and “gastrulation” (Figure 5G, adjusted p < 0.05). Collectively, this indicates that loss of the P2-proximal CTCF binding site caused a delay in in vitro hematopoietic differentiation, providing functional support for a mild decrease in P2-derived *Runx1* transcription at or prior to the mesoderm stage.

**Figure 5.**
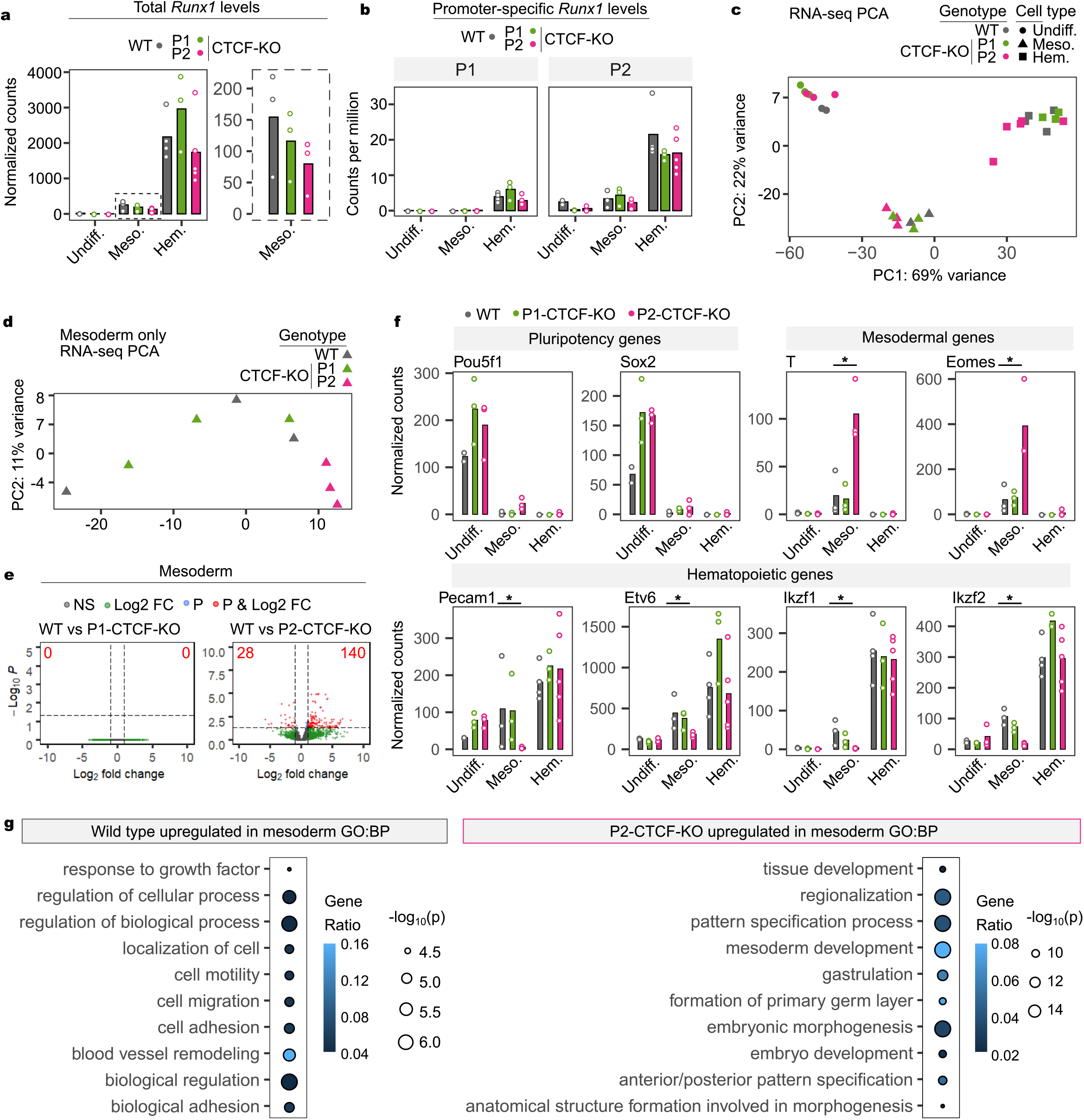
*Runx1* spatiotemporal expression is slightly altered after loss of P2-proximal CTCF. **A)** Total *Runx1* levels in poly(A)-minus RNA-seq in the cell types and genotypes indicated. The expanded graph with a dashed outline shows data just for mesodermal cells on a different axis. **B)** Promoter-specific *Runx1* levels for each promoter in the cell types and genotypes indicated. **C)** PCA of all poly(A)-minus RNA-seq replicates. **D)** PCA of mesoderm RNA-seq samples. **E)** Volcano plots showing differentially expressed genes (DEGs, adjusted p<0.05, fold change>1) in P2-CTCF-KO compared to wild type mesoderm. **F)** Expression of lineage marker genes across differentiation in the genotypes indicated (*, adjusted p<0.05, fold change>1). **G)** GO term biological processes associated with the DEG list between wild type and P2-CTCF-KO mesoderm. Gene ratios and -log_10_ p values are indicated.

## Discussion

In the present study we used a cutting-edge 3C-based method to reveal dynamic changes in the *Runx1* chromatin architecture in four-dimensions, i.e. the three-dimensional folding of chromatin over developmental time. Tiled-C (Oudelaar et al. (2020) analysis of the *Runx1* regulatory domain provided an unprecedented high-resolution view of *Runx1* chromatin architecture during *in vitro* differentiation from mESC through Flk1^+^ mesoderm to differentiating HPCs. Our detailed dissection of *Runx1* transcriptional regulation during developmental hematopoiesis shed new light on regulatory mechanisms of complex large developmental genes. We found that *Runx1* resides in a preformed, transcription-independent and evolutionarily conserved main TAD that is present throughout differentiation. Within this TAD dynamic sub-structures formed over development, namely sub-TADs spanning the *Runx1* gene that appeared specifically in HPCs. We showed that promoter-proximal CTCF sites played a role in the maintenance of *Runx1* sub-TADs, but, interestingly, not in mediating the dynamic changes in E-P interactions associated with hematopoietic differentiation. Yet, loss of the P2-proximal CTCF site led to delayed hematopoietic differentiation and disrupted gene expression specifically at the Flk1^+^ mesoderm stage, possibly by slight reductions in *Runx1* levels. Finally, we found that during hematopoietic development from mesoderm to HPCs, the *Runx1* promoters switched from interacting with enhancers located throughout the TAD, including the gene desert, to primarily interacting with cis-elements closer to the gene and within the tissue-specific sub-TADs. This refines the region within which functional enhancers important for driving hematopoietic-specific *Runx1* expression are likely to be found. These hematopoietic enhancers may represent novel therapeutic targets in leukemia, similar to what was recently shown for the *RUNX1* +23 enhancer (Mill et al. 2019). The 4D regulatory interactions at *Runx1* described here were missed in previous reports (Chen et al. 2019, Marsman et al. 2017, Wilson et al. 2016) as those lacked the required high-resolution, nor included a developmental time series.

In line with TADs at other developmentally regulated loci (Brown et al. 2018, Hug et al. 2017, Oudelaar et al. 2020, Paliou et al. 2019), the overall 1.1 Mb *Runx1* TAD formed prior to differentiation and in the absence of virtually any gene transcription. The mechanism behind its establishment is likely CTCF/cohesin-mediated loop extrusion (Alipour and Marko 2012, Fudenberg et al. 2016, Nasmyth 2001, Sanborn et al. 2015) as a predominant convergence of CTCF motifs was observed at the main *Runx1* TAD boundaries, similar to what was found at other TADs (de Wit et al. 2015, Rao S. S. et al. 2014, Vietri Rudan et al. 2015). In addition to these preformed chromatin structures, increasing CTCF-CTCF interactions were observed upon *Runx1* activation that might reflect a higher rate or processivity of loop extrusion in the *Runx1* TAD. This was recently also observed for α-globin (Oudelaar et al. 2020), implicating that common mechanisms underlie chromatin architecture changes during transcriptional activation at different sized gene loci. However, we also observed differences in chromatin structures between smaller and larger genes, in that two tissue-specific sub-TADs formed over differentiation, leading to a sub-compartmentalization of *Runx1*. Sub-compartmentalization is not generally seen at smaller genes, such as α-globin, which reside entirely within one tissue-specific sub-TAD (Hanssen et al. 2017, Hay et al. 2016). This difference might simply reflect the smaller size of the α-globin domain compared to the larger size of the *Runx1* regulatory domain (97 kb compared to 1.1 Mb, respectively). The sub-TAD encompassing the entire α-globin regulatory unit was shown to represent a discrete functional unit that delimits enhancer activity (Hanssen et al. 2017). We did not see evidence for this in Runx1 as E-P interactions were not significantly changed upon sub-TAD perturbation by promoter-proximal CTCF sites. The finding that the *Runx1* tissue specific sub-TADs were strongest in HPCs, which express high levels of *Runx1*, suggests they may be similar to the gene-body-associated domains (GADs) recently observed at genes highly expressed in hematopoietic cells (Zhang C. et al. 2020). The formation of sub-TADs/GADs within the gene-body of actively transcribed genes suggests that these structures, instead of being caused by a CTCF-dependent mechanism like loop extrusion, may instead be dependent on transcriptional processes (Zhang C. et al. 2020). Indeed, our data indicates that, as reported for other promoters (Cho et al. 2018, Harrold et al. 2020, Schwessinger et al. 2020), the *Runx1* promoters may function as chromatin boundaries in a CTCF-independent manner, which would explain the residual sub-TADs observed in P1/P2-CTCF-KO cells. Together, our findings suggest that loop extrusion and transcription-related mechanisms may act in concert to produce dynamic chromatin structures during differentiation.

Interestingly, deletion of the *Runx1* P2 promoter-proximal CTCF binding site resulted in a developmental delay in hematopoietic mesoderm as shown by increased expression of early pluripotency and mesodermal markers at the expense of later hematopoietic ones. Although studies in cell lines indicate that CTCF sites are required for TAD formation (Nora et al. 2017, Rao et al. 2017, Schwarzer et al. 2017, Wutz et al. 2017), interestingly, promoter-proximal CTCF sites have been suggested to play a role in E-P interactions and gene transcription (Canzio et al. 2019, Kubo et al. 2021, Lee et al. 2017, Ren et al. 2017, Schuijers et al. 2018, Zhou et al. 2021). Therefore, a plausible explanation for the observed differentiation delay could be that promoter-proximal CTCF loss leads to a later or perturbed onset of *Runx1* expression, as Runx1 is well known to promote hematopoietic commitment (Swiers et al. 2013). Although *Runx1* expression in the total mesodermal cell population was not significantly altered, a decreased trend was seen. This surprisingly robust *Runx1* expression upon promoter-proximal CTCF-site loss may reflect the residual sub-TADs still found which could be attributed to redundant CTCF sites not targeted in this study (redundancy between CTCF sites has been observed before (Hanssen et al. 2017, Kentepozidou et al. 2020, Schwessinger et al. 2020)), or the *Runx1* promoters themselves. An alternative explanation could be the relatively asynchronous development of cells in culture, where some cells may have had enough time to restore *Runx1* expression levels. Finally, compared to in vitro differentiation, promoter-proximal CTCF-site loss may be more detrimental *in vivo*, where *Runx1* levels are subject to tight spatiotemporal control and changes to *Runx1* levels or dosage lead to knock-on effects on differentiation timing (Cai et al. 2000, Lacaud et al. 2004, Lie et al. 2018, Song et al. 1999, Wang Q. et al. 1996). Together this indicates that even subtle changes in *Runx1* levels, such as the trend seen in P2-CTCF-KO mesoderm, have the potential to alter hematopoietic developmental dynamics. This underlines that *Runx1* requires an exceptionally fine-scale spatiotemporal transcriptional control and isolation from neighboring regulatory domains to support its pivotal role in development. Given that *Runx1* has important functions in development and human disease (de Bruijn and Dzierzak 2017, Levanon and Groner 2004, Mevel et al. 2019), an increased understanding of dynamic cis-regulatory mechanisms underpinning its regulation will be vital to future efforts to develop potential therapeutic approaches to manipulate *RUNX1* expression in human blood disorders.

## Materials and methods

### Cell culture

E14-TG2a mESCs (Handyside et al. 1989) were cultured in GMEM medium supplemented with 100 mM non-essential amino acids, 100 mM sodium pyruvate, 10% FCS, 2 mM L-glutamine, 100 *µ*M β-mercaptoethanol (all Gibco), and 1% Leukemia Inhibitory Factor (prepared in house). Cells were passaged using 0.05% trypsin (Gibco) every 2-3 days. The 416B mouse immortalized myeloid progenitor cell line (Dexter et al. 1979) was cultured at 2-8×10^5^ cells/ml in Fischer’s medium with 20% horse serum and 2 mM L-glutamine (all Gibco).

### Hematopoietic differentiation of mESCs

Differentiation of mESCs was performed using a modified serum-free protocol (Pearson et al. 2015, Sroczynska et al. 2009b) in StemPro-34 (SP34, Gibco) supplemented with 40X defined serum replacement, 2 mM L-glutamine (Gibco), and 0.5 mM ascorbic acid, 0.45 mM monothioglycerol (Sigma). mESCs were seeded at a density of 5 x10^4^ cells/ml into SP34 medium plus BMP-4 (R&D, 5 ng/ml). At day 3, bFGF and Activin A were added (R&D, 5 ng/ml). At day 4, single cell suspension was generated from embryoid bodies using 0.05% trypsin. FACS-isolated Flk1+ mesodermal cells were cultured at 5 x10^4^ cells/cm^2^ in SP34 plus SCF (Peprotech) and VEGF (R&D), 10 ng/ml each. After a further 3 days of culture, adherent and suspension cells were treated with 0.05% trypsin and analyzed.

### Flow cytometry and cell sorting

Cells were stained in PBS plus 10% FCS with the antibodies listed in Supp. Table 3. Dead cells were identified with Hoechst 33258. Cells were analyzed using a Fusion 2 (Becton Dickinson) flow cytometer.

### Colony forming unit assays

Unsorted cells at day 6 (4+2) of differentiation were cultured in MethoCult 04434 (Stem Cell Technologies) in 35 mm dishes. Colonies were counted after 10 days.

### Immunocytochemistry and confocal microscopy

Cells were plated into glass bottom 24-well plates (ibidi) and fixed after culture for 10 mins with 4% paraformaldehyde (Sigma), permeabilized, and labeled using antibodies (Supp. Table 3) for 1 hour at room temperature. Imaging was performed using a Zeiss 880 laser scanning confocal microscope.

### Deletion of CTCF sites using CRISPR/Cas9

Single guide RNAs (sgRNAs, Supp. Table 5) were designed (crispr.mit.edu) to flank conserved CTCF motifs. sgRNAs were cloned into pSpCas9(BB)-2A-Puro V2.0 (Ran et al. 2013) containing two sgRNAs and confirmed by sequencing. mESCs were transfected with 5 *µ*g plasmid using Lipofectamine 2000 (Invitrogen) and puromycin selected (1 *µ*g/ml). Single colonies were isolated by limiting dilution (Gruzdev et al. 2019). To detect larger on-target deletions (Mianne et al. 2017, Owens et al. 2019, Teboul et al. 2020), PCR amplification was done (500 bp to 5 kb, Supp. Table 5). Sequencing confirmed the presence of two distinct deletion alleles and the retention of DpnII restriction sites.

### Copy counting by droplet digital PCR (ddPCR)

To further rule out the presence of undesired complex genotypes (Kosicki et al. 2018), copy counting was performed across the targeted regions by droplet digital PCR (ddPCR) as described (Owens et al. 2019). Reactions were performed in duplex, amplifying from an internal control and a test region in every reaction. Internal control was located on mouse chromosome 4 (not targeted in these experiments and karyotypically stable in mESCs (Codner et al. 2016)). Test regions were amplified directly over the targeted region to detect loss of allele (LOA) and at 100 bp and 1 kb up- and down-stream from sgRNA target sites (Supp. Table 5)(Owens et al. 2019). Reactions (22 *µ*l) contained 11 *µ*l QX200 ddPCR EvaGreen Supermix (Bio-Rad), 25-50 ng genomic DNA purified using DNeasy Blood and Tissue Kit (Qiagen), 250 nM each of internal control primer, and 125 nM each of test primer. Standard reagents and consumables supplied by Bio-Rad were used. Ratios between test and internal control amplicons was determined in QuantaSoft software (Bio-Rad). Ratios were normalized to the mean ratio of three test amplicons located on different non-targeted chromosomes (1, 6, and 7) to determine relative copy numbers of the test amplicons.

### Chromatin interaction analysis (Tiled-C)

Tiled-C was performed on between 7.7×10^4^-1×10^6^ cells using a low-input protocol described previously (Oudelaar et al. 2017a, Oudelaar et al. 2017b, Oudelaar et al. 2020). Cells were crosslinked using 2% formaldehyde for 10 minutes. DpnII (NEB) digestion was performed shaking overnight at 37 °C. Ligation was performed overnight at 16 °C. Samples were treated with RNAse A (Roche) for 30 minutes at 37 °C and decrosslinked using Proteinase K (Thermo Fisher) overnight at 65 °C. Digestion efficiency was quantified by qPCR (Supp. Table 5). Libraries with greater than 70% digestion efficiency were used (mean 83%). Up to 1 *µ*g DNA was sonicated using a Covaris ultrasonicator. End-repair, adaptor ligation, and PCR addition of indices (7-11 cycles) was done using NEBNext Ultra II DNA library prep kit (NEB). Biotinylated capture probes 70 nt in length were designed against every DpnII restriction fragment in a 2.5 Mb window centered on *Runx1* (chr16:91,566,000-94,101,999). Sequences were BLAT-filtered and synthesized in-house (Oudelaar et al. 2020). A pooled capture reaction was performed on 1 *µ*g of each indexed 3C library. Washing of captured material was done using Nimblegen SeqCap EZ hybridisation and wash kit (Roche) and captured sequences were isolated using M-270 Streptavidin Dynabeads (Invitrogen). PCR amplification was performed for 12 cycles. Amplified DNA was purified and a second capture and PCR amplification step were performed. Libraries were sequenced on two Illumina NextSeq high-output 150 cycle runs (paired end).

### Analysis of Tiled-C data

Tiled Capture-C data was processed at 2 kb resolution as described (Oudelaar et al. 2020). Fastqs were analyzed using the CCSeqBasic CM5 pipeline (Telenius Jelena M. et al. 2020) (https://github.com/Hughes-Genome-Group/CCseqBasicF/releases). Individual samples were analyzed before merging biological replicates. PCR duplicated-filtered bam files were converted to sam files (samtools) and then into sparse raw contact matrices (Tiled_sam2rawmatrix.pl, https://github.com/oudelaar/TiledC). ICE normalization was done using HiC-Pro (2.11.1)(Imakaev et al. 2012, Servant et al. 2015) and matrices were imported into R (3.6.0). Matrices were plotted (TiledC_matrix_visualisation.py, https://github.com/oudelaar/TiledC) with a threshold between the 90^th^ - 95^th^ percentile. PCA was done on log normalized counts (DESeq2,1.24.0) (Love et al. 2014). Merged ICE normalized contact matrices were scaled to the mean number of total interactions (14631865) across samples. Virtual Capture-C plots were generated by sub-setting the matrices on individual viewpoints of interest. E-P contacts were quantified in count and ICE-normalized matrices from the viewpoint of the bin containing each promoter and bins overlapping previously published *Runx1* enhancers (Supp. Table 2). TADs were detected by visual inspection. Intra-TAD interactions were calculated by quantifying the ratio between intra- and extra-TAD interactions for each bin within the TAD in each sample. TAD boundary contacts were quantified between bins overlapping the four outermost CTCF sites at each of the centromeric and telomeric ends of the main TAD.

### Analysis of Hi-C data

Publicly available Hi-C data in mESCs (Bonev et al. 2017) was analyzed as previously described (Oudelaar et al. 2020). Data were analyzed using HiC-Pro (Servant et al. 2015) with ICE normalization (Imakaev et al. 2012), and plotted using python as described above.

### DeepC prediction of chromatin architecture

DeepC predictions were performed as described (Schwessinger et al. 2020). Briefly, deepC was trained using a transfer learning approach on distance stratified and percentile binned Hi-C data from mESCs (Bonev et al. 2017), withholding mouse chromosome 16 (that contains *Runx1*) and chromosome 17 from training. Training was done in two stages. First, a convolutional neural network was trained to predict chromatin features given a 1 kb DNA sequence input. The chromatin features cover open chromatin, transcription factor binding, including CTCF, and histone modifications using publicly available DNase-seq, ATAC-seq and ChIP-seq peaks across a range of cell types. Second, a neural network using a convolutional module followed by a dilated convolutional module was trained to predict Hi-C data given 1 Mb of DNA sequence input. The convolutional filters of the first network are used in the transfer learning to seed the filters of the convolutional module. Promoter-proximal CTCF site deletions were modelled by mutating the region spanning CRISPR/Cas9 deletion alleles that were confirmed by sequencing and predicting the chromatin interactions of the reference and deletion alleles.

### Gene expression analysis (RNA-seq)

RNA was isolated from 1×10^3^-2.5×10^6^ cells using QIAzol (Qiagen). Total RNA was extracted using miRNeasy Mini kit (Qiagen). RNA integrity was determined using a 4200 TapeStation RNA ScreenTape (Agilent). Ribosomal RNA was depleted from 2.5 *µ*g total RNA per sample of undifferentiated mESC and 416B cells using the RiboMinus™ Eukaryote System v2 (Invitrogen). A poly A selection module (NEB) was used to extract poly A minus RNA and was eluted directly in First Strand Synthesis Reaction Buffer. cDNA was synthesised using the NEBNext Ultra directional library prep kit. Adapter ligation and 8-15 cycles of PCR was performed. Libraries were sequenced on Illumina NextSeq high-output 75 cycle kit (paired-end).

### RNA-seq analysis

Fastq files were mapped to the mouse genome (mm9) using STAR (2.6.1d) (Dobin et al. 2013). PCR duplicates were removed using picard-tools (2.3.0) MarkDuplicates. Counts per million (CPM)-normalised bigwig files were generated using deeptools (3.0.1) (Ramirez et al. 2016). A blacklist file was used to exclude mapping artifacts. Bigwig files were converted to bedGraph (ucsctools (373)) and imported into R. Mean CPM was calculated for each merged sample. Reads were assigned using subread (2.0.0) featureCounts (Liao et al. 2014). For poly(A)-minus RNA-seq data reads were assigned to both exons and introns. Assigned counts were imported into R and analyzed using DESeq2 (1.24.0) (Love et al. 2014). Sample clustering was performed on log normalized counts. Differential expression analysis was done using DESeq2 (adjusted p value > 0.05, fold change > 1). Volcano plots were made using EnhancedVolcano (1.2.0) (Blighe 2019). GO terms were calculated using goseq (1.36.0)(Young et al. 2010) and KEGG.db (3.2.3) (Carlson 2016). Gene expression was visualized using plotCounts and ggplot2 (3.3.0)(Wickham 2016). Promoter-specific counts were quantified from over a 5kb window downstream of each TSS (Runx1-P1 chr16:92823811-92828811; Runx1-P2 chr16:92695073-92700073) using bedtools (2.25.0)(Quinlan and Hall 2010).

### Chromatin accessibility analysis (ATAC-seq and DNaseI-seq)

ATAC-seq libraries were generated in differentiated E14-TG2a-RV mESCs (stably transfected with a Venus reporter at the 3′ end of *Runx1* (Harland et al. 2021) and a hsp68-mCherry-Runx1+23 enhancer-reporter transgene in the *Col1a1* locus). Libraries were generated as previously described (Buenrostro et al. 2013). 2-5×10^4^ differentiated cells were FACS-isolated, resuspended in cold lysis buffer and incubated for 10 mins on ice. Cells were centrifuged, supernatant discarded and resuspended in 10 *µ*l transposition mix. Samples were incubated for 30 minutes at 37 °C and quenched using 1.1 *µ*l 500 mM EDTA. Reactions were centrifuged and incubated at 50 °C for 10 mins. A total of thirteen cycles of PCR were performed as in Buenrostro et al. (2013) with transposition reaction as a template. PCR reactions were purified using MinElute PCR purification kit (Qiagen). Libraries were sequenced using Illumina NextSeq 75 cycle kit with paired-end reads. Fastq files were mapped to the mouse genome (mm9) (NGseqBasic VS2.0)(Telenius Jelena and Hughes 2018). PCR duplicate-filtered bam files from individual samples were merged and filtered to remove reads mapping to chrM, ploidy regions, or the *Runx1*-Venus targeting construct (chr16:92602138-92605899, chr16:92606403-92609879), and only reads with short (<100bp) insert sizes were retained. RPKM-normalised bigwig files were generated using deeptools (3.0.1) (Ramirez et al. 2016). DNaseI-seq data in undifferentiated mESCs were downloaded from GEO (GSM1014154) (Vierstra et al. 2014) and analyzed as ATAC-seq data were. Peaks were called using MACS2 (Liu 2014) (p-value > 0.05).

### CTCF binding (ChIP-seq)

CTCF ChIP was conducted using Millipore ChIP agarose kit (Millipore). 1×10^6^ crosslinked 416B cells were lysed and sonicated using a Covaris ultrasonicator. Sonicated chromatin was diluted using dilution buffer and 50 *µ*L was removed as the 5% input control. 2uL CTCF antibody (Supp. Table 3) was added to 1 mL chromatin and incubated overnight at 4 °C. Decrosslinking was done at 65 °C overnight. DNA was purified using phenol-chloroform-isoamylalcohol (25:24:1, Sigma) and enrichment was determined using qPCR (Supp. Table 5). NEBNext Ultra II DNA Library Prep Kit (NEB) with 11 cycles of PCR was used to prepare sequencing libraries. CTCF ChIP libraries were sequenced using Illumina NextSeq high-output 75 cycle kit (paired-end).

### CTCF ChIP-seq analysis and de novo CTCF motif annotation

CTCF ChIP-seq was performed in 416B cells and publicly available E14 mESC data (Handoko et al. 2011) was downloaded from GEO (GSE28247). Fastq files were mapped to the mouse genome (mm9) (NGseqBasic VS2.0)(Telenius Jelena and Hughes 2018). De novo CTCF motifs were identified in CTCF ChIP-seq data using meme (4.9.1_1)(Bailey and Elkan 1994) as described previously (Hanssen et al. 2017). CTCF peaks were called using MACS2 (Zhang Y. et al. 2008) with parameters -p 0.02 using input track as a control. 2000 peaks were sampled using bedtools (2.25.0)(Quinlan and Hall 2010) and flanking regions were extracted from the sampled peaks. Sequences of sampled peaks and flanking regions were retrieved and a background file was generated using fasta-get-markov -m 0. A de novo motif file was generated using meme with options -revcomp -dna -nmotifs 1 -w 20 -maxsize 1000000 - mod zoops. De novo motifs were identified in CTCF peaks using fimo with options -motif 1 -thresh 1e-3.

### Targeted bisulfite sequencing

DNA methylation analysis was performed as previously described (Jeziorska et al. 2017). Genomic DNA (gDNA) was extracted from 1-5×10^6^ cells using DNeasy Blood and Tissue Kit (Qiagen) and 250 ng gDNA, or Universal Methylated Mouse DNA Standard (Zymo Research) was bisulfite converted using EZ DNA Methylation-GoldTM Kit (Zymo Research). Nested PCR primer sets (Supp. Table 5) were designed to amplify 281-379 bp overlapping target regions. External PCR reactions were performed on 1*µ*L bisulfite converted DNA using HotStarTaq DNA Polymerase (Qiagen). Internal nested PCR reactions were performed using 1*µ*L of the external PCR reaction. Amplicons were size selected by gel and purified. 250 ng equimolar PCR amplicons were combined for each biological sample and indexed using NEBNext Ultra II DNA Library Prep Kit for Illumina (NEB) with 6 PCR cycles. Reads were quality and adapter trimmed using trim galore/0.3.1 (https://github.com/FelixKrueger/TrimGalore). Reads were mapped to an in silico bisulfite converted genome using bismark/0.20.0 (Krueger and Andrews 2011). Percentages of methylated CpG dinucleotides were determined using bismark methylation extractor. Bedgraph output files were filtered on CpG dinucleotides with coverage greater than 100 reads and imported into R. Average methylated CpG dinucleotide percentages were plotted over each region using ggplot2.

### Statistical tests

All statistical tests were performed in R. Tiled-C contact data were non-normal (Shapiro-Wilks test, P < 2×10^−16^) and so non-parametric Kruskal-Wallis test with Dunn’s post-hoc comparisons test was applied. Post-hoc testing with Dunn’s test was applied when Kruskal-Wallis test was significant and p-values were adjusted using the Holm method. A significance threshold of p < 0.05 was used for all statistical tests.

## Data availability

Sequencing data will be made available upon publication.

## Acknowledgements

We thank Christina Rode, Emanuele Azzoni, Vincent Frontera, Ruth Williams and Tatjana Sauka-Spengler for helpful discussions and advice. We thank Nick Crump and Tom Milne for expert technical assistance and advice. We thank Kevin Clark, Sally Clark, and Paul Sopp from the WIMM FACS facility for assistance with cell sorting and technical expertise. This work was supported by programmes in the MRC Molecular Hematology Unit Core award to M.F.T.R.d.B (MC_UU_12009/2) and J.R.Hu (MC_UU_00016/14), MRC MR/N00969X/1 and Wellcome Trust (106130/Z/14/Z) to J.R.Hu, Wellcome Trust Doctoral Programmes supported A.C. (108870/Z/15/Z), R.S. (203728/Z/16/Z) and A.M.O. (105281/Z/14/Z), who was also supported by the Stevenson Junior Research Fellowship (University College, Oxford), an EPSRC studentship supported J.R.Ha, a Clarendon Scholarship L.G, and S.d.O was supported by an MRC Project Award (MR/N00969X/1) to J.R.Hu. The WIMM Flow Cytometry facility is supported by the MRC HIU, MRC MHU [MC_UU_12009], NIHR Oxford BRC and John Fell Fund [131/030 and 101/517], EPA fund [CF182 and CF170], WIMM Strategic Alliance awards [G0902418 and MC_UU_12025]. The Wolfson Imaging Centre Oxford is supported by the Medical Research Council via the WIMM Strategic Alliance (G0902418), the Molecular Haematology Unit (MC_UU_12009), the Human Immunology Unit (MC_UU_12010), the Wolfson Foundation (Grant 18272), and by an MRC/BBSRC/EPSRC grant (MR/K015777X/1) to MICA—NanoscopyOxford (NanO): Novel Super-resolution Imaging Applied to Biomedical Sciences, Micron (107457/Z/15Z).

## Author Contributions

D.D.G.O, A.M.O., D.J.D., J.R.Hu., M.F.T.R.d.B. designed and conceptualized the study. D.D.G.O., G.A., A.M.O., D.J.D., A.C., J.R.Ha., A.B., performed experiments. D.D.G.O., A.M.O., J.R.Ha., R.S., D.J., performed data analysis and interpretation. L.G., S.d.O, D.J., generated vital reagents. D.D.G.O, G.A., A.M.O., D.J.D., A.C., J.R.Hu., and M.F.T.R.d.B wrote the paper. All authors approved of the final version to be published.

