## Supplemental data for "Dynamic *Runx1* chromatin boundaries affect gene expression in hematopoietic development"

#### Supplementary Figure legends

**Supplementary Figure 1*. In vitro* hematopoietic differentiation mimics endothelial-to-hematopoietic transition *in vivo*. A)** Confocal imaging (maximum intensity projections) of immunocytochemical (ICC) staining of day 7 cultures with CD31 shown in white, Runx1 in red, CD41 in green, and DAPI in blue. A colony of HE cells and emerging progenitors is outlined by the yellow dashed line. A Runx1+CD31+CD41-/lo HE cell is indicated by the solid white arrowhead. Emerging hematopoietic progenitors are indicated by the hollow white arrowhead. **B)** May-Grünwald staining of cytospins of FACS-isolated CD41+ CD45- cells. **C)** Hematopoietic colony formation assays (CFU-C) on day 6 whole cultures. The number of colonies formed per 10,000 cells seeded is shown.

**Supplementary Figure 2*.* Tiled-C matrices of individual replicates in wild type cells over hematopoietic differentiation.** Tiled-C matrices from individual experiments are shown at 2 kb resolution and are total count and ICE normalised. All matrices are visualized with a threshold set at the 94$^{th}$ percentile.

**Supplementary Figure 3*.* CTCF binding close to *Runx1* promoters and promoters genome wide is associated with promoter activity. A)** *Runx1* locus showing chromatin marks and poly(A)-minus RNA-seq in undifferentiated mESCs and 416B cells. Public data that were reanalyzed include mESC and 416B DNaseI (Vierstra et al. 2014) and mESC CTCF (Handoko et al. 2011). The orientation of CTCF motifs are indicated underneath peaks. **B)** Analysis of *Runx1* promoter methylation by targeted bisulfite sequencing of undifferentiated mESCs, hematopoietic 416B cells. *In vitro* methylated DNA and an additional negative control site are shown as controls. **C)** Enrichment of CTCF peaks in undifferentiated mESCs and 416B cells at genomic features annotated using HOMER (4.7) annotatePeaks.pl. **D)** Association between CTCF binding within 5 kb of transcription start sites (TSS) shown for active and inactive TSSs in undifferentiated mESCs and 416B hematopoietic cells. CTCF ChIP-seq was reanalyzed from previously published data (Handoko et al. 2011). *, chi-square test, $p < 1x10^{-10}$.

**Supplementary Figure 4*.* DeepC predictions of Runx1 chromatin interactions and promoter-proximal CTCF site deletions in undifferentiated mESCs. A)** Tiled-C (top panel) and deepC prediction of chromatin interactions (bottom panel) at the *Runx1* locus in undifferentiated mESCs. The overall 1.1Mb *Runx1* TAD indicated below the matrices was substantially concordant between the deepC predictions and real Tiled-C data. CTCF occupancy (Handoko et al. 2011) and motif orientation are indicated. **B, C)** Tiled-C (top panels) and deepC predictions (bottom panels) of chromatin interactions after P2-CTCF (B) and P1-CTCF (C) deletion. DeepC predicted that, in undifferentiated mESCs, P2-CTCF deletion would lead to increased interactions across the boundary (B, bottom panel, pink upside down triangle with its tip at the P2-CTCF site) and a decrease in a stripe of interactions emanating from the CTCF site (B, bottom panel, dark grey line at 45$^{o}$). Both features are visible in Tiled-C data in undifferentiated mESCs (B, top panel). Little change in interactions was seen after loss of P1-CTCF in mESC (C, top panel), which was similar to the prediction by deepC (C, bottom panel). CTCF binding (Handoko et al. 2011) and motif orientation are indicated below the matrices.

**Supplementary Figure 5*.* Strategy for the generation of P1-CTCF knock-out clones using CRISPR/Cas9. A)** Schematic of the *Runx1* P1 promoter and upstream promoter-proximal CTCF site with single guide RNAs (sgRNAs), short-range (SR), medium range (MR), longer range 3 kb (LR-3kb) and longer range 5 kb (LR-5kb) PCR primer locations indicated. The *de novo* CTCF motif identified under the CTCF peak (P1-CTCF) is indicated with its orientation. Droplet digital PCR (ddPCR) amplicon positions are indicated and were spaced upstream (5') or downstream (3') of sgRNA cut positions by approximately the indicated distances. One ddPCR amplicon was placed between sgRNA cut sites to detect loss of the CTCF motif (LOA). Poly(A) minus RNA-seq and CTCF ChIP-seq tracks for 416B cells are included. Publicly available DNaseI (Vierstra et al. 2014) and H3K27ac ChIP-seq (Schutte et al. 2016) data for 416B cells were reanalyzed and shown in the respective tracks. Evolutionary sequence conservation shown is vertebrate conservation by PhastCons (Siepel et al. 2005). **B)** Close-up view of dual sgRNA (2xsgRNA) targeting strategy to delete a conserved CTCF motif using CRISPR/Cas9. The location of the nearest DpnII restriction sites is indicated and were not disrupted by targeting. **C)** Gel electrophoresis images of PCR samples from six independent sub-clones derived from two separate targeting events. **D)** Base calls from sequencing of sub-clones revealing homozygous deletion of the CTCF motif. **E)** Copy number analysis by ddPCR across the targeted region and on three diploid (2n) control chromosomes.

**Supplementary Figure 6. Strategy for the generation of P2-CTCF knock-out clones using CRISPR/Cas9. A)** Schematic of the *Runx1* P2 promoter and adjacent upstream promoter-proximal CTCF site with single guide RNAs (sgRNAs), short-range (SR), medium range (MR), longer-range 3 kb (LR-3kb) and longer range-5 kb (LR-5kb) PCR primer locations indicated. The *de novo* CTCF motif identified under the CTCF peak (P2-CTCF) is indicated with its orientation. Droplet digital PCR (ddPCR) amplicon positions are indicated and were spaced upstream (5') or downstream (3') of sgRNA cut positions by approximately the indicated distances. One ddPCR amplicon was placed in between sgRNA cut sites to detect loss of the CTCF motif (LOA). Poly-A minus RNA-seq and CTCF tracks for 416B cells are included. Publicly available DNaseI (Vierstra et al. 2014) and H3K27ac ChIP-seq (Schutte et al. 2016) data for 416B cells were reanalyzed and shown in the respective tracks. CpG island annotations are from UCSC genome browser (Kent et al. 2002). Evolutionary sequence conservation shown is vertebrate conservation by PhastCons (Siepel et al. 2005). **B)** Close-up view of dual sgRNA (2xsgRNA) targeting strategy to delete a conserved CTCF motif using CRISPR/Cas9. The location of the nearest DpnII restriction sites is indicated and were not disrupted by targeting. **C)** Gel electrophoresis images of PCR samples from six independent sub-clones derived from two separate targeting events. **D)** Base calls from sequencing of sub-clones revealing homozygous deletion of the CTCF motif. **E)** Copy number analysis by ddPCR across the targeted region and on three diploid (2n) control chromosomes.

**Supplementary Figure 7. Loss of CTCF binding at P2-CTCF in P2-CTCF-KO clones.** CTCF ChIP-seq in six undifferentiated mESC clones (three P1-CTCF-KO and three P2-CTCF-KO). CTCF occupancy is shown at the *Runx1* P2 promoter and a positive control gene promoter that also binds CTCF. Binding at *Runx1* P1 could not be assessed as binding is too low in undifferentiated mESCs.

**Supplementary Figure 8. CTCF binding upstream of *Runx1* P1 or P2 promoter is not required for hematopoietic differentiation of mESCs.** Flow cytometry analysis of wild type, P1-CTCF-KO and P2-CTCF-KO cells at differentiation day 4 (Flk1) and day 7 (Ter119, CD41 and CD45). Relative cell numbers (percentages of live cells or Terr119- cells) and absolute cell numbers are shown for each genotype or cell type. Results are from three independent clones per genotype and 10 independent experiments (n=7-11).

**Supplementary Figure 9. Tiled-C matrices of individual replicates in P1-CTCF-KO cells over hematopoietic differentiation. A)** Individual matrices are shown at 2 kb resolution and are total count and ICE normalised. All matrices are visualized with a threshold set at the 94$^{th}$ percentile. **B)** Principal component analysis (PCA) of Tiled-C data for individual replicates, with each cell type shown in a different shape and each genotype colored differently.

**Supplementary Figure 10. Tiled-C matrices of individual replicates in P2-CTCF-KO cells over hematopoietic differentiation. A)** Individual matrices are shown at 2 kb resolution and are total count and ICE normalised. All matrices are visualized with a threshold set at the 94$^{th}$ percentile. **B)** Principal component analysis (PCA) of Tiled-C data for individual replicates, with each cell type shown in a different shape and each genotype colored differently.

**Supplementary Figure 11. Enhancer-promoter contacts are maintained across hematopoietic differentiation despite loss of CTCF binding and disrupted *cis*-interactions.** Enhancer-promoter contacts quantified between *Runx1* P1 or P2 promoters and all previously published hematopoietic enhancers (Supp. Table 2). The median contacts and interquartile ranges are shown. There were no significant differences between genotypes for any of the enhancer-promoter or in any of the cell types examined (Kruskal-Wallis and Dunn's test, adjusted p = 1.0).

**Supplementary Figure 12. Enhancer-promoter interactions are maintained despite perturbed chromatin architecture. A)** Normalized Tiled-C matrix in wild type HPCs showing *Runx1* tissue-specific sub-TADs. *Runx1* promoters and previously identified enhancers are indicated. RNA-seq and ATAC-seq in wild type HPCs is shown, alongside CTCF in 416B cells. Virtual Capture-C (obtained from Tiled-C data, see methods) from the viewpoint of the +23 enhancer in HPCs is shown for the genotypes indicated. The dark color corresponds to the mean interactions in four independent replicates, while the lighter color above shows the standard deviation. **B)** Total enhancer-promoter contacts are shown for *Runx1* P1 and P2 promoters in the different cell types and for the genotypes indicated. Each bar represents median interaction count and error bars show the interquartile range.

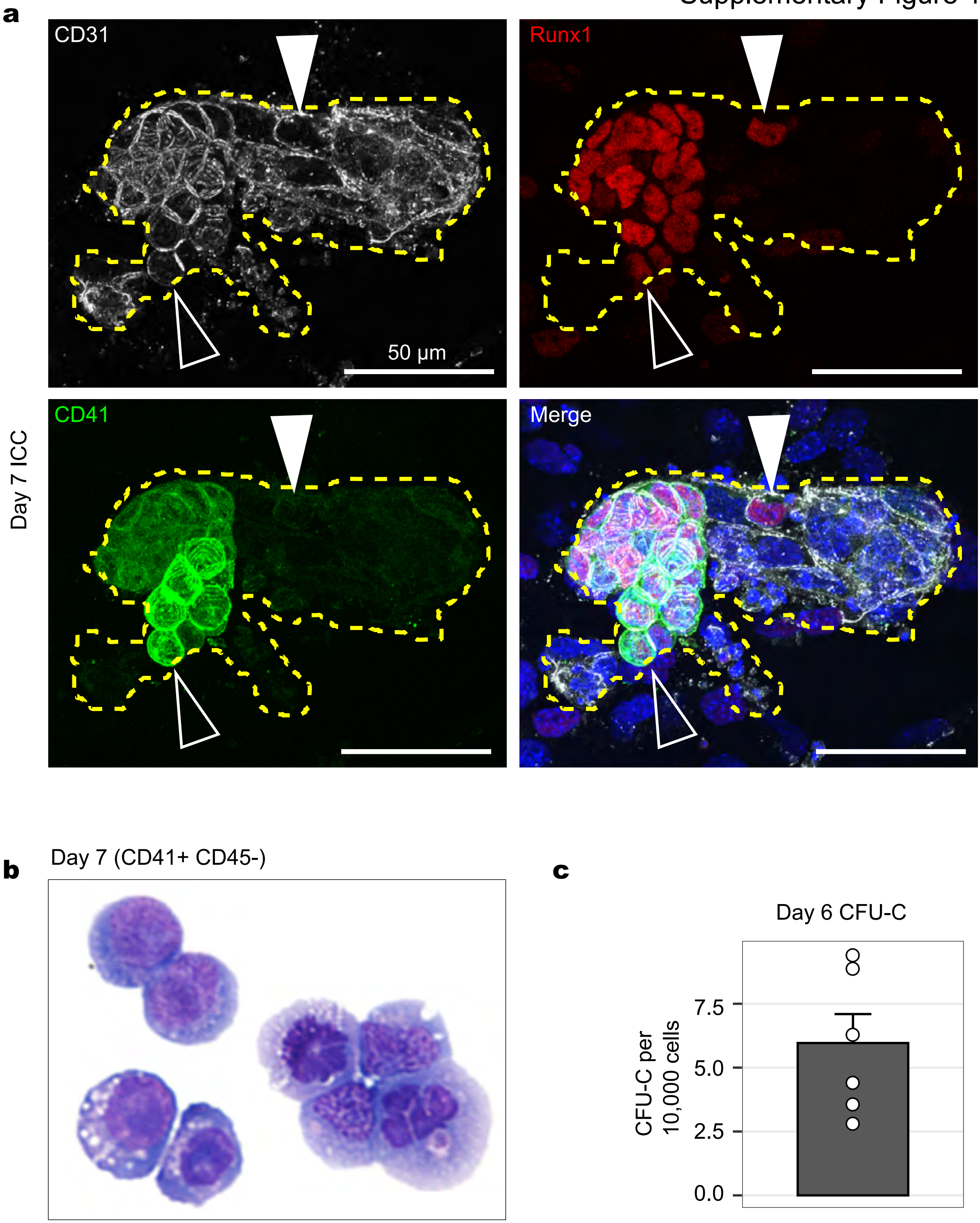
a
CD31
Runx1
50 µm
CD41
Merge
Day 7 ICC
b
Day 7 (CD41+ CD45-)
c
Day 6 CFU-C
CFU-C per 10,000 cells
7.5
5.0
2.5
0.0

Supplementary Figure 2

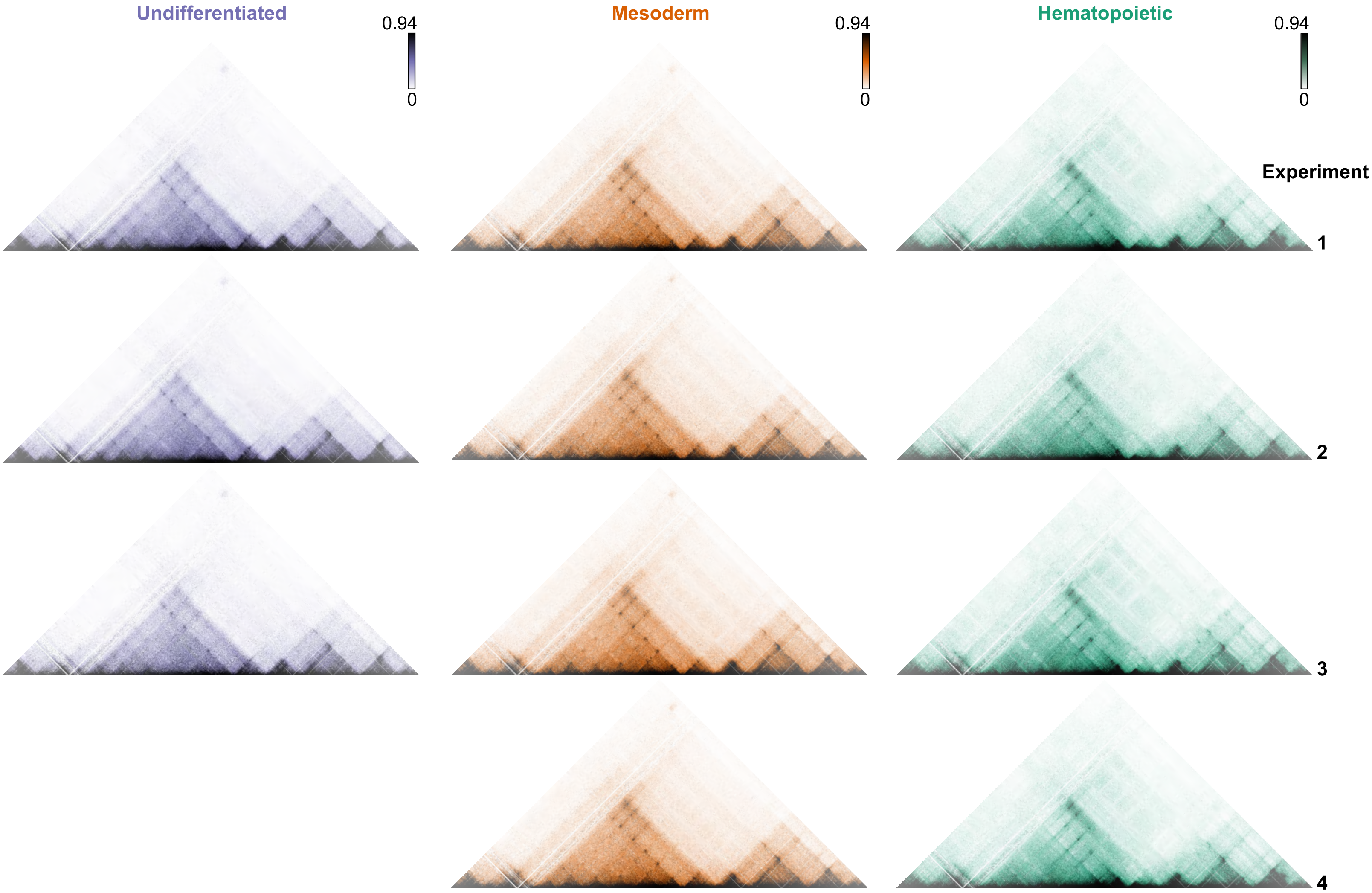

### Supplementary Figure 3

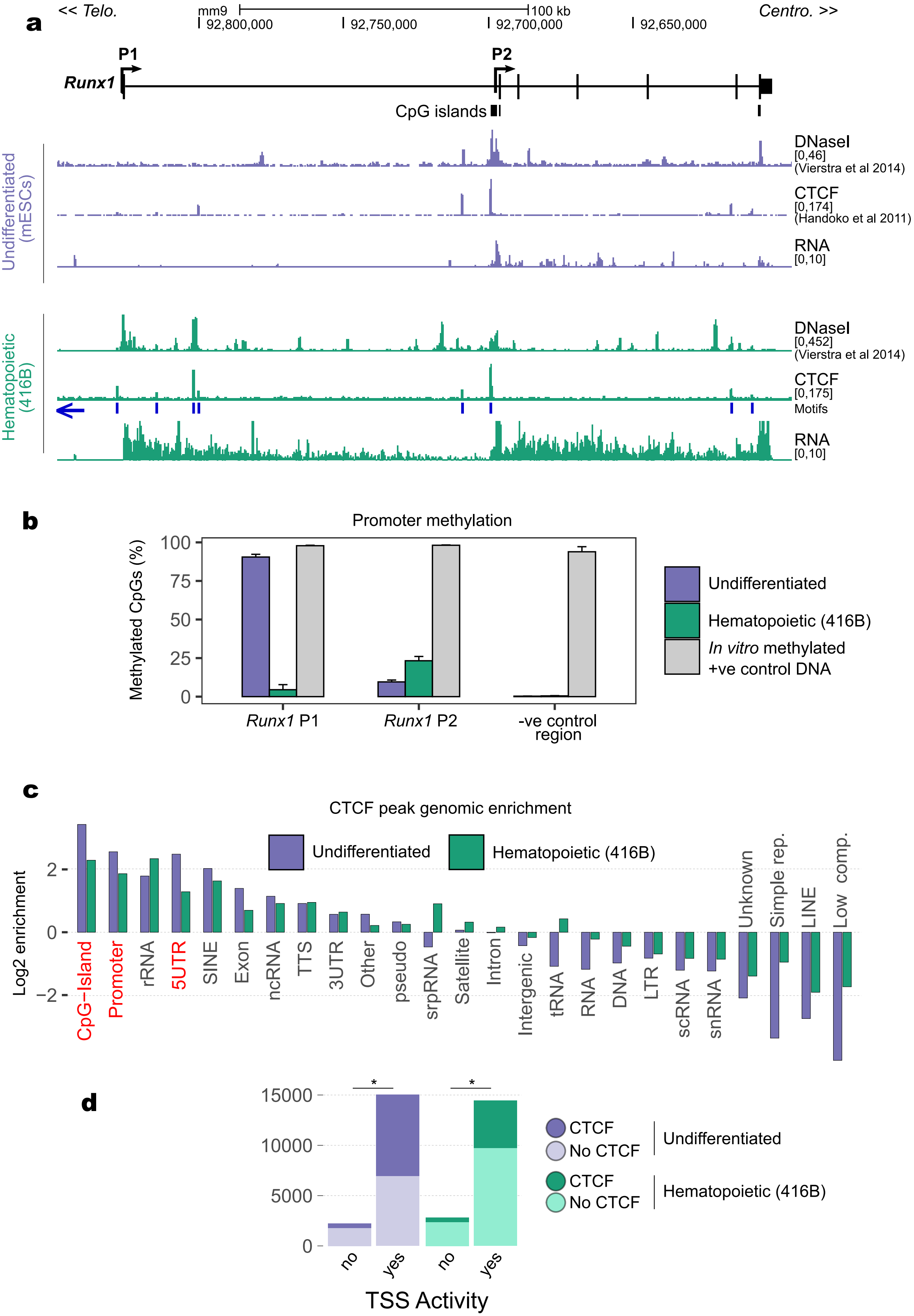

### Supplementary Figure 4

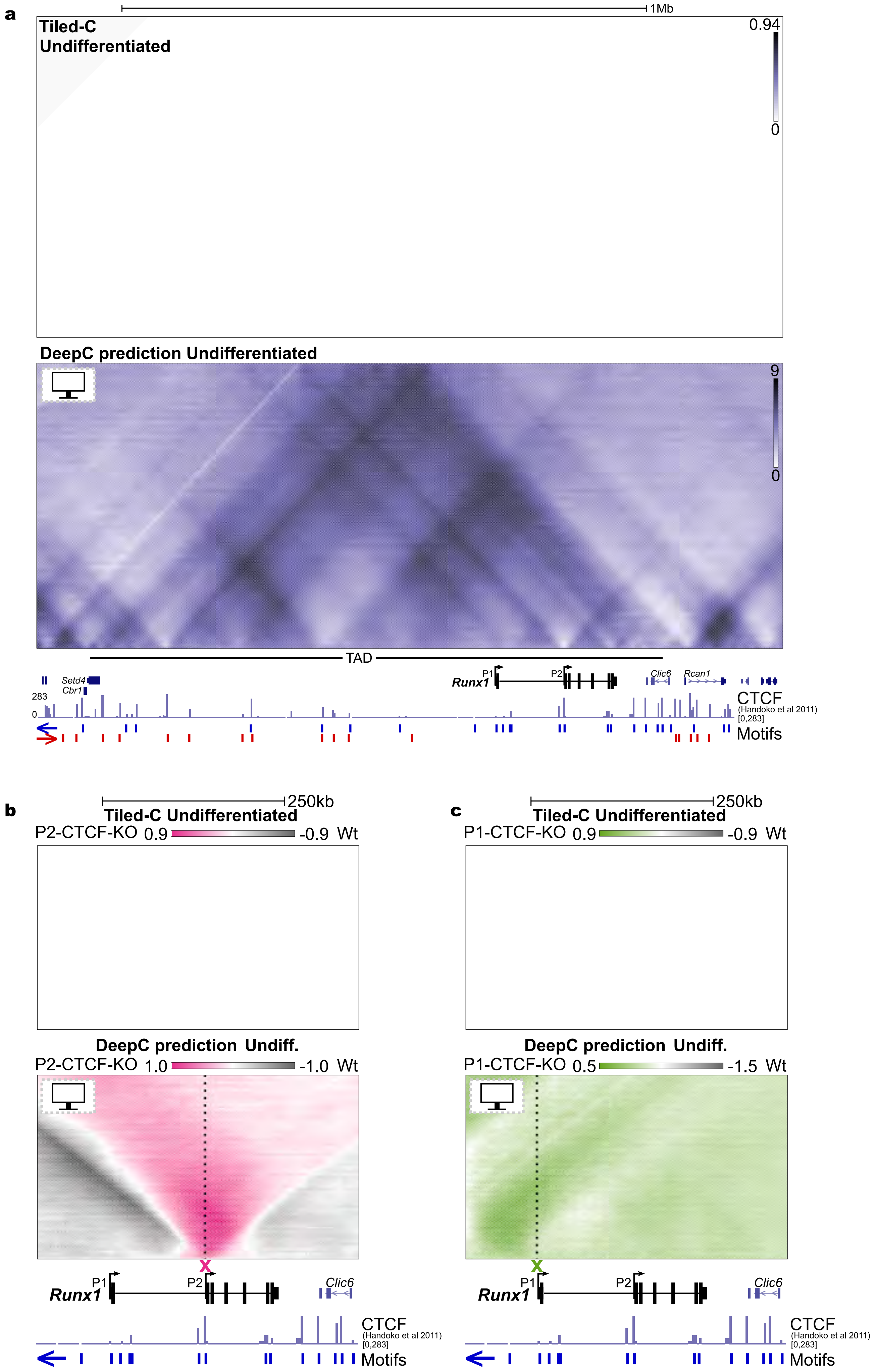

Supplementary Figure 5

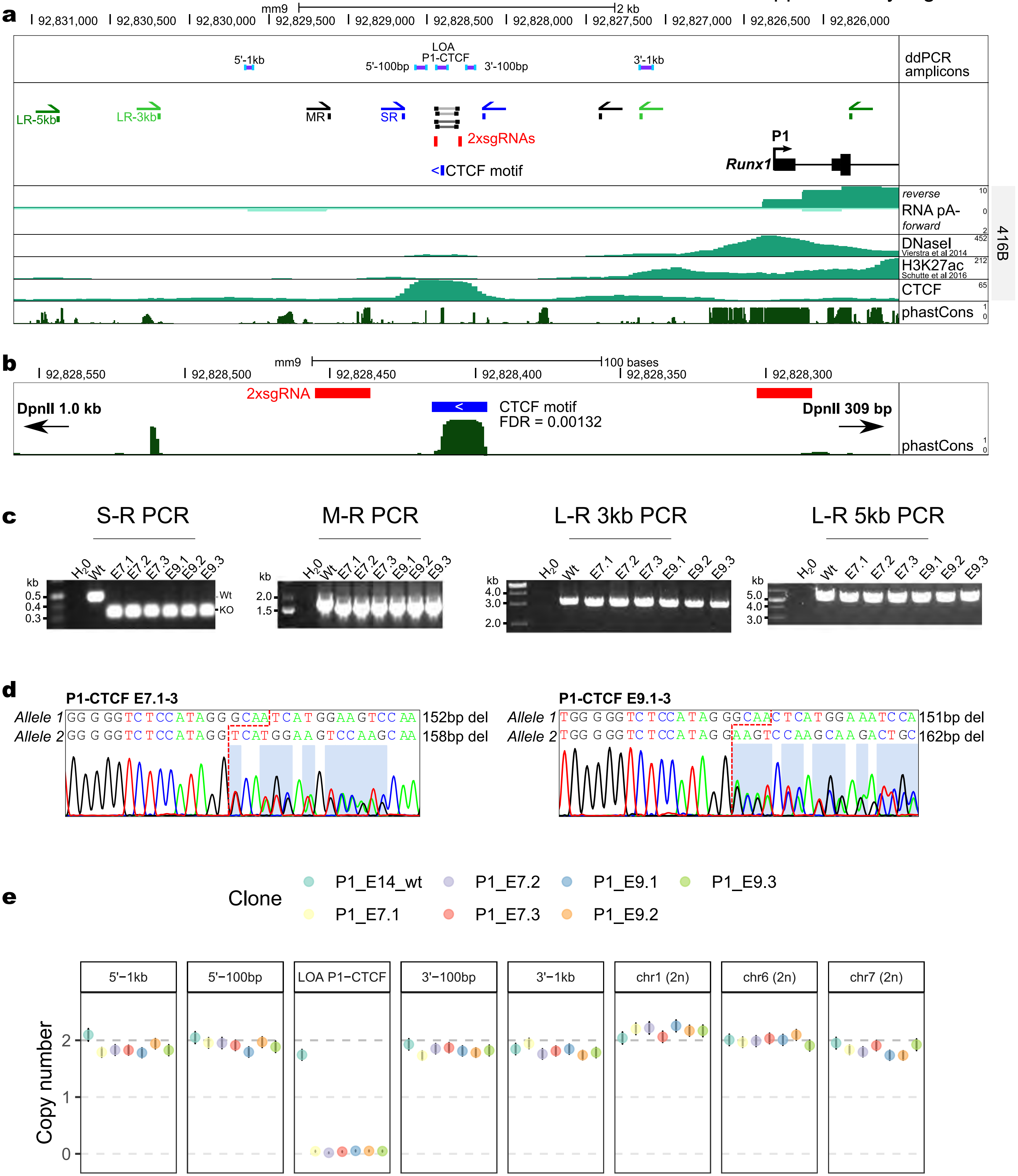

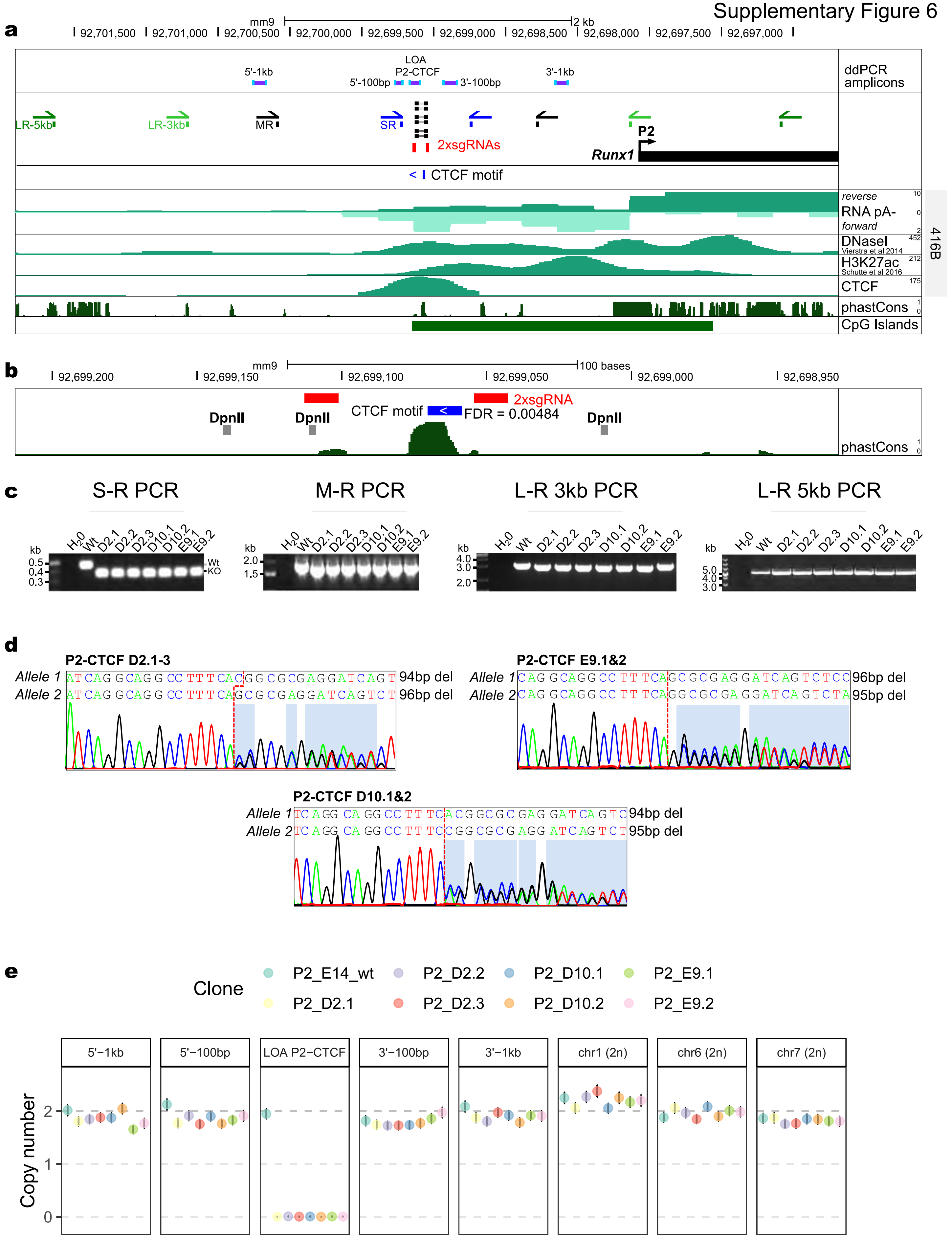
a
mm9
2 kb
92,701,500
92,701,000
92,700,500
92,700,000
92,699,500
92,699,000
92,698,500
92,698,000
92,697,500
92,697,000
5'-1kb
LOA P2-CTCF
5'-100bp
3'-100bp
3'-1kb
ddPCR amplicons
LR-5kb
LR-3kb
MR
SR
2xsgRNAs
P2
Runx1
CTCF motif
reverse
RNA pA-
forward
DNaseI
Vierstra et al 2014
H3K27ac
Schutte et al 2016
CTCF
phastCons
CpG Islands
416B
b
mm9
100 bases
92,699,200
92,699,150
92,699,100
92,699,050
92,699,000
92,698,950
2xsgRNA
DpnII
CTCF motif
FDR = 0.00484
phastCons
c
S-R PCR
M-R PCR
L-R 3kb PCR
L-R 5kb PCR
H2O
Wt
D2.1
D2.2
D2.3
D10.1
D10.2
E9.1
E9.2
kb
Wt
KO
d
P2-CTCF D2.1-3
Allele 1
Allele 2
94bp del
96bp del
P2-CTCF E9.1&2
96bp del
95bp del
P2-CTCF D10.1&2
94bp del
95bp del
e
Clone
P2_E14_wt
P2_D2.1
P2_D2.2
P2_D2.3
P2_D10.1
P2_D10.2
P2_E9.1
P2_E9.2
5'−1kb
5'−100bp
LOA P2−CTCF
3'−100bp
3'−1kb
chr1 (2n)
chr6 (2n)
chr7 (2n)
Copy number

### Supplementary Figure 7

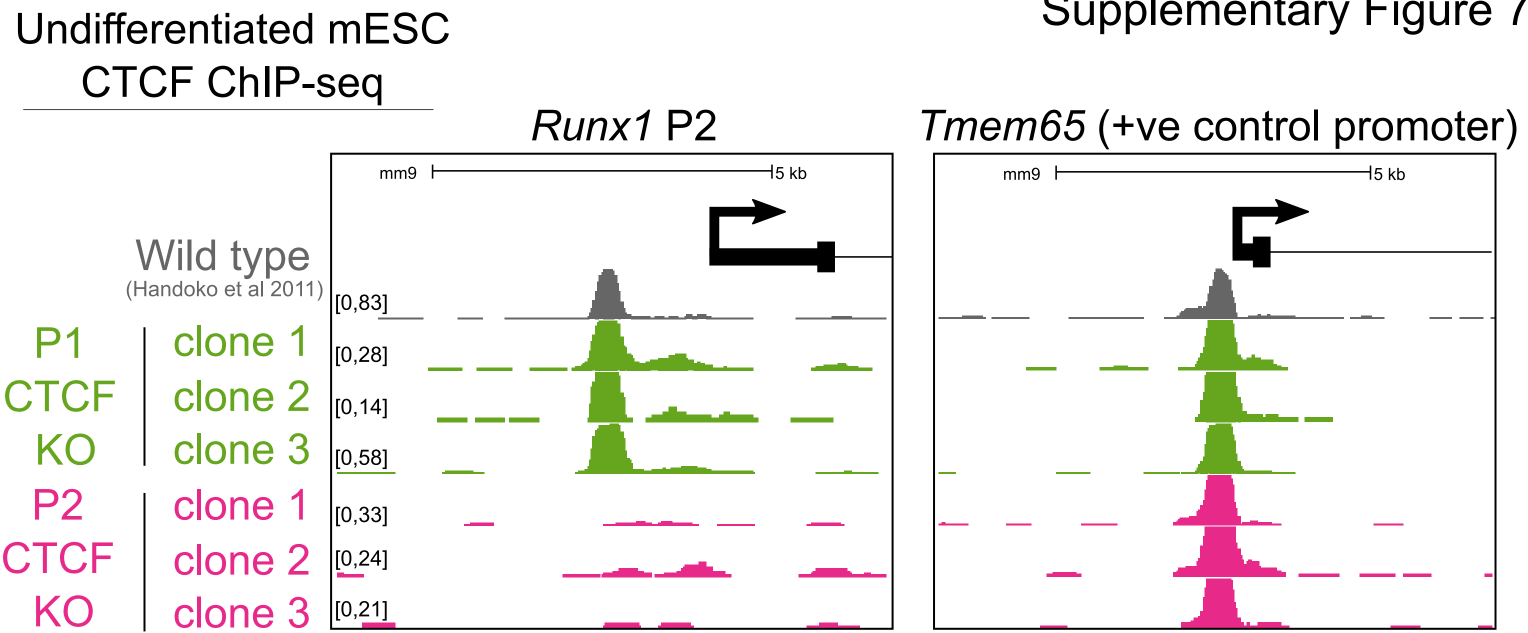

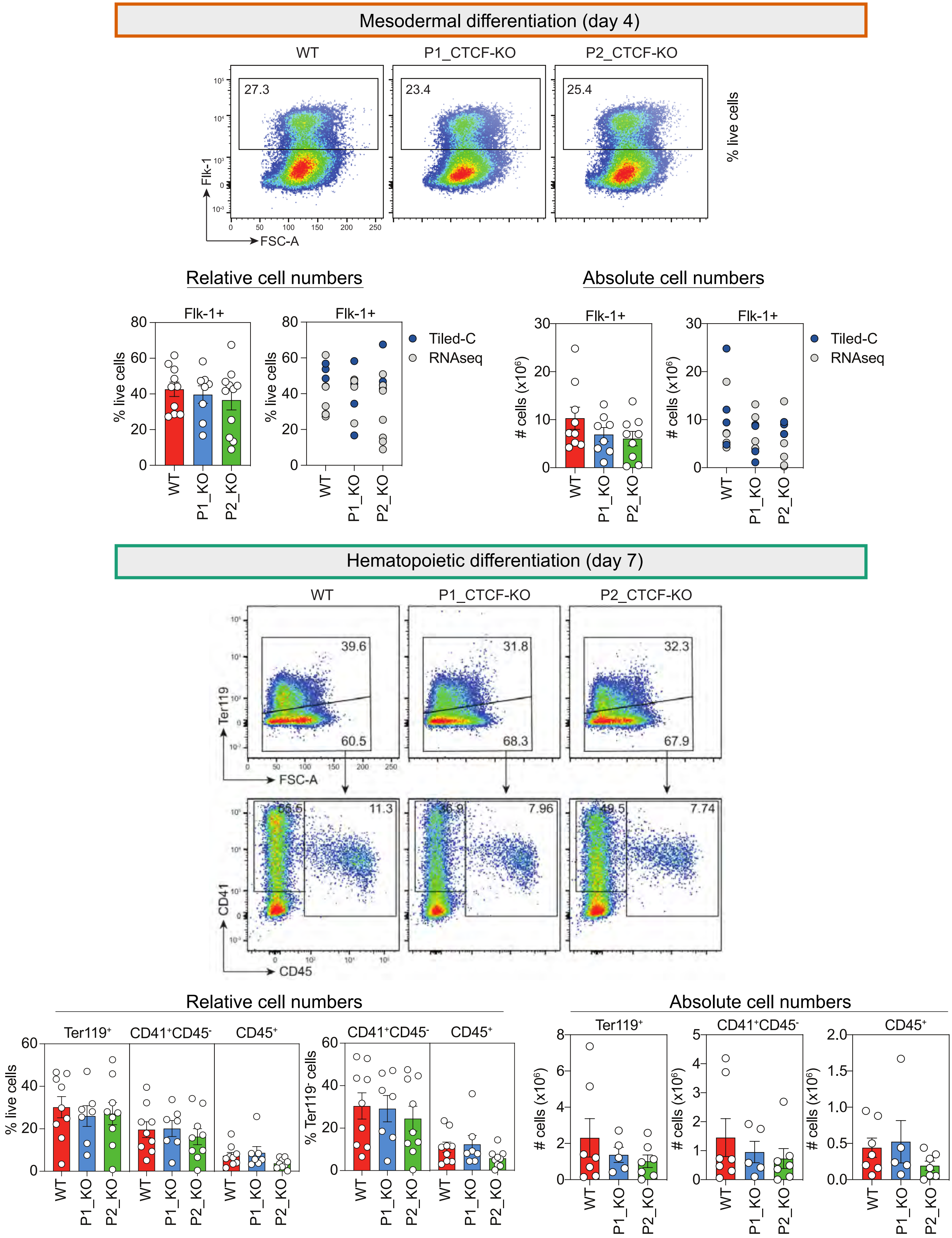

Mesodermal differentiation (day 4)
WT
P1_CTCF-KO
P2_CTCF-KO
27.3
23.4
25.4
Flk-1
FSC-A
% live cells
Relative cell numbers
Absolute cell numbers
Flk-1+
% live cells
### cells (x10^6)
Tiled-C
RNAseq
WT
P1_KO
P2_KO
Hematopoietic differentiation (day 7)
39.6
60.5
31.8
68.3
32.3
67.9
Ter119
FSC-A
11.3
7.96
7.74
CD41
CD45
Relative cell numbers
Absolute cell numbers
Ter119+
CD41+CD45-
CD45+
% live cells
% Ter119- cells
### cells (x10^6)

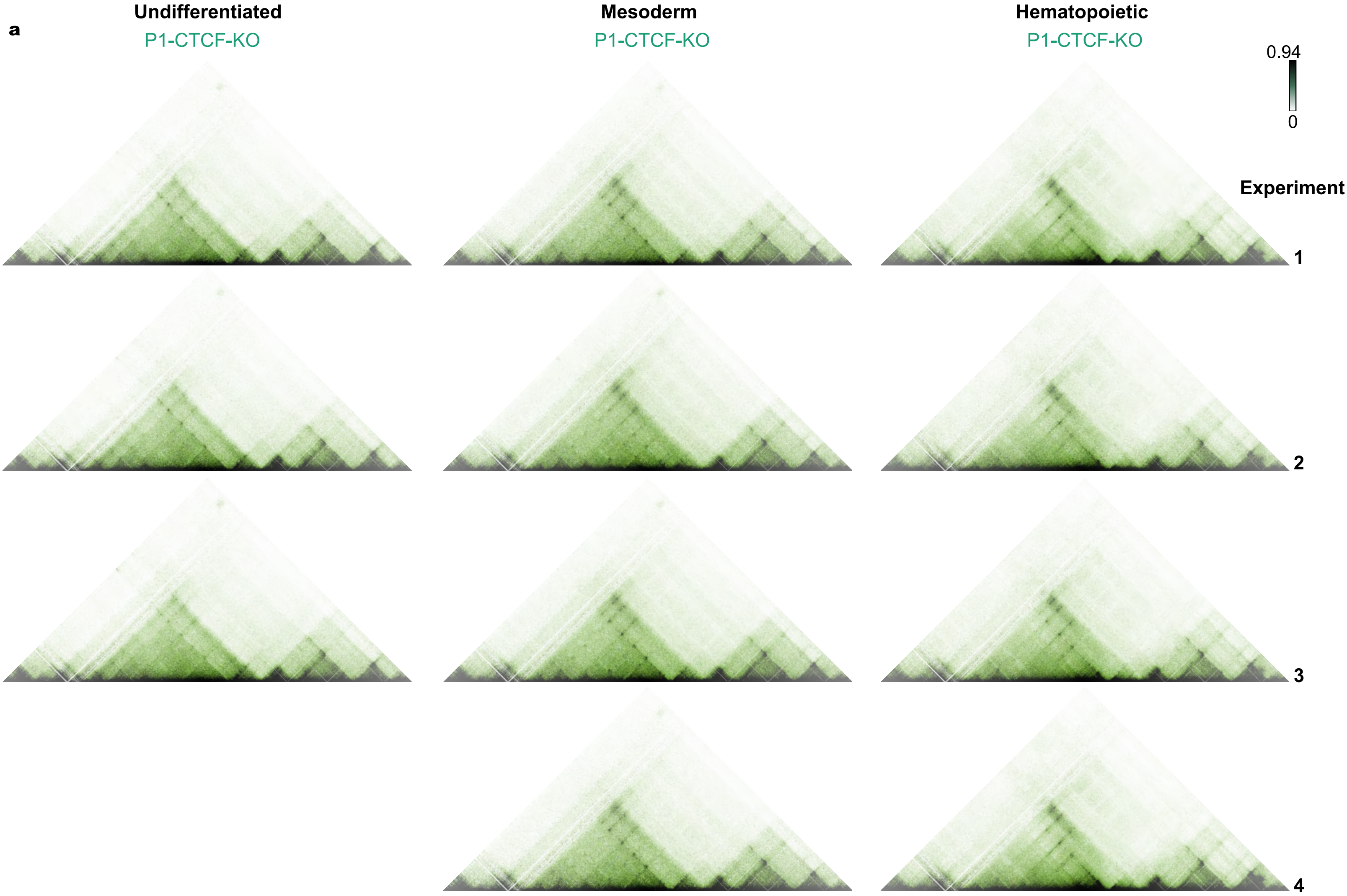
a
Undifferentiated
P1-CTCF-KO
Mesoderm
P1-CTCF-KO
Hematopoietic
P1-CTCF-KO
0.94
0
Experiment
1
2
3
4

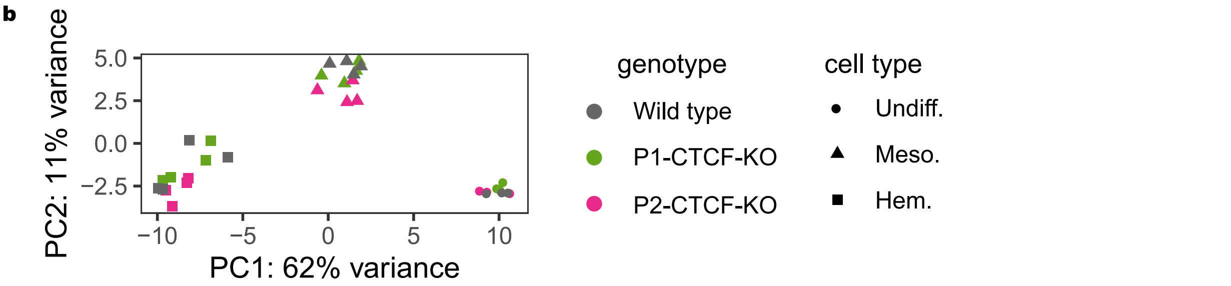
b
PC2: 11% variance
5.0
2.5
0.0
−2.5
−10
−5
0
5
10
PC1: 62% variance
genotype
Wild type
P1-CTCF-KO
P2-CTCF-KO
cell type
Undiff.
Meso.
Hem.

Supplementary Figure 10

**a**

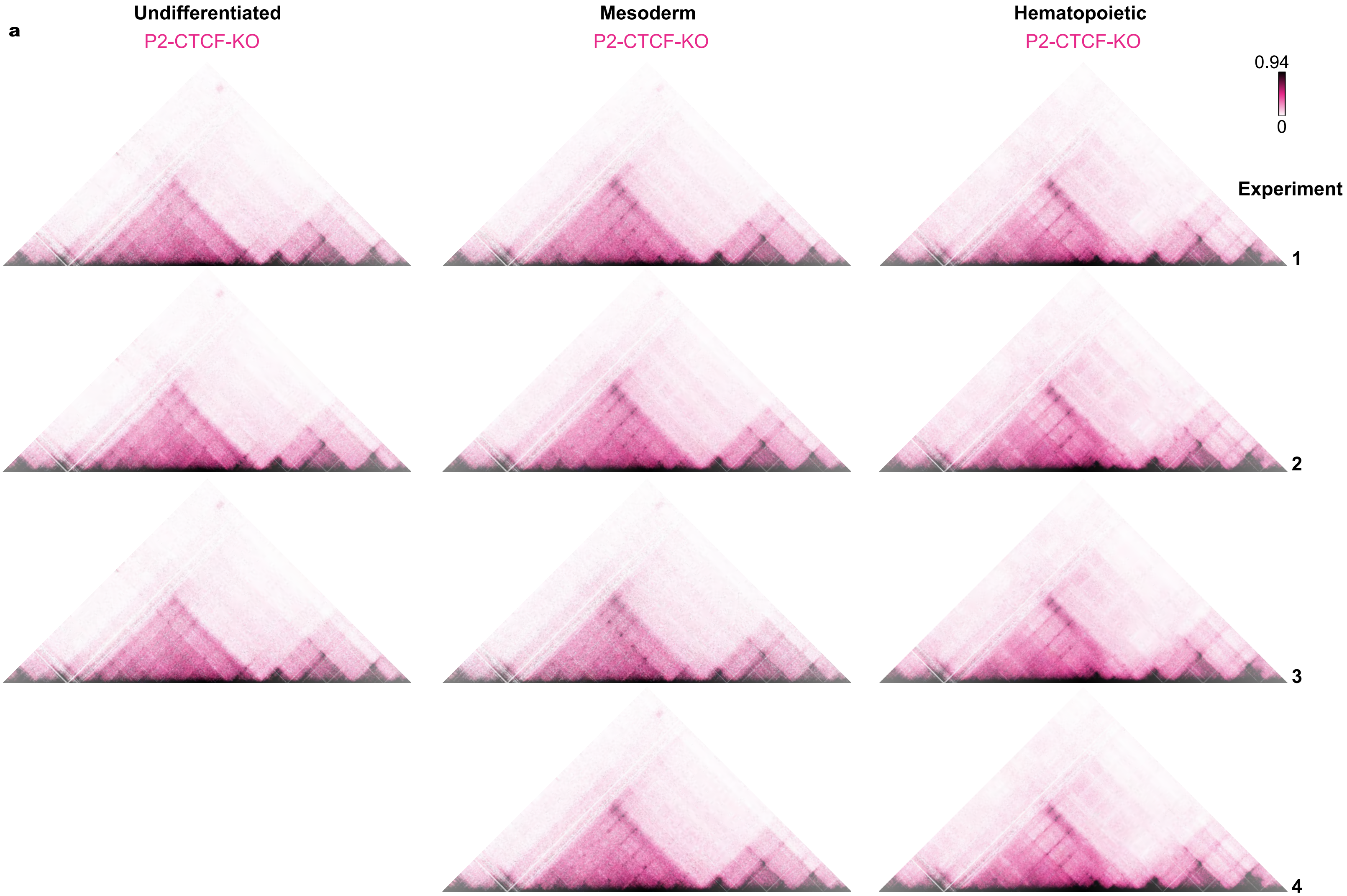


**b**

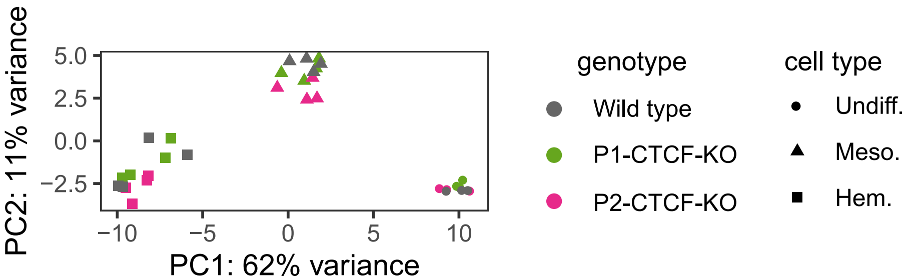

**a**

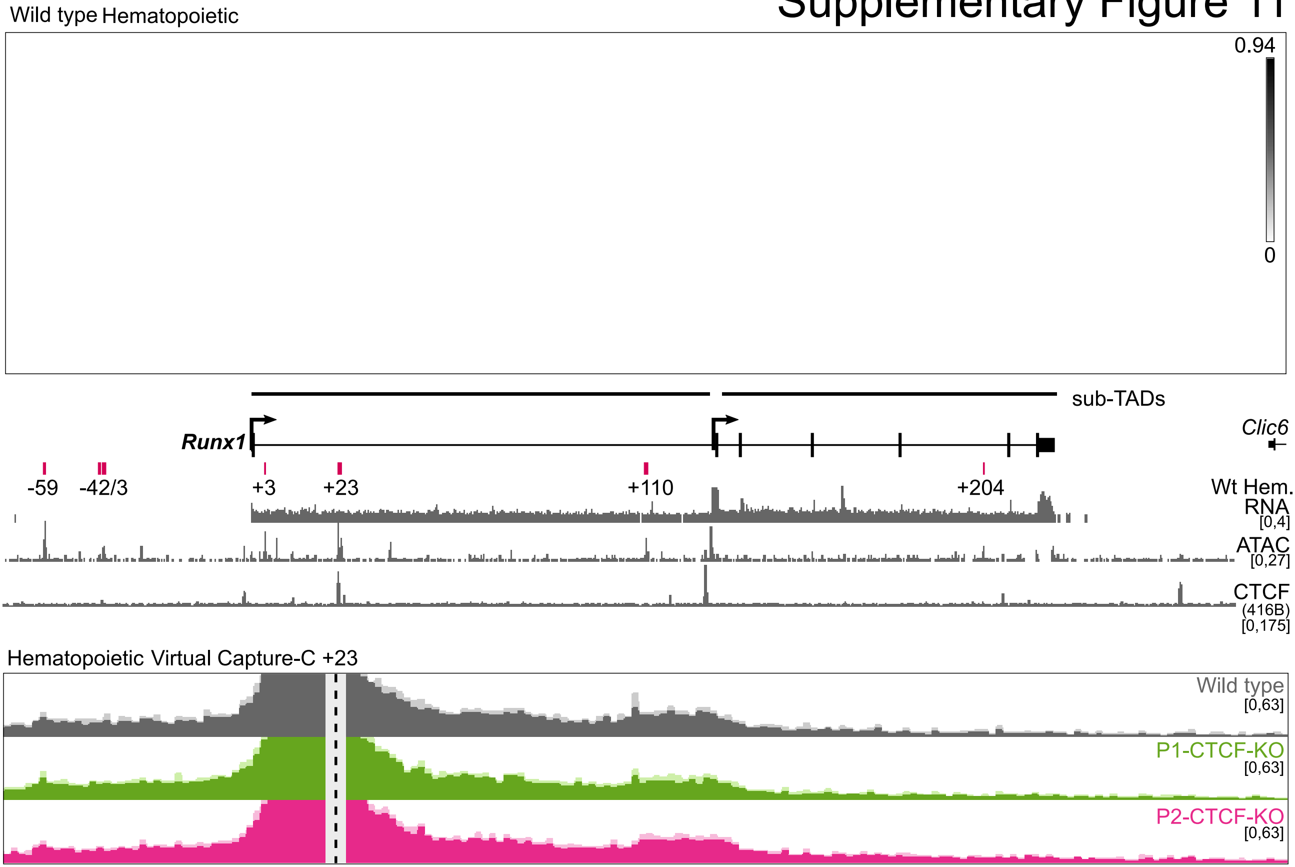
Wild type Hematopoietic
0.94
0
sub-TADs
Runx1
Clic6
-59
-42/3
+3
+23
+110
+204
Wt Hem. RNA [0,4]
ATAC [0,27]
CTCF (416B) [0,175]
Hematopoietic Virtual Capture-C +23
Wild type [0,63]
P1-CTCF-KO [0,63]
P2-CTCF-KO [0,63]

**b**

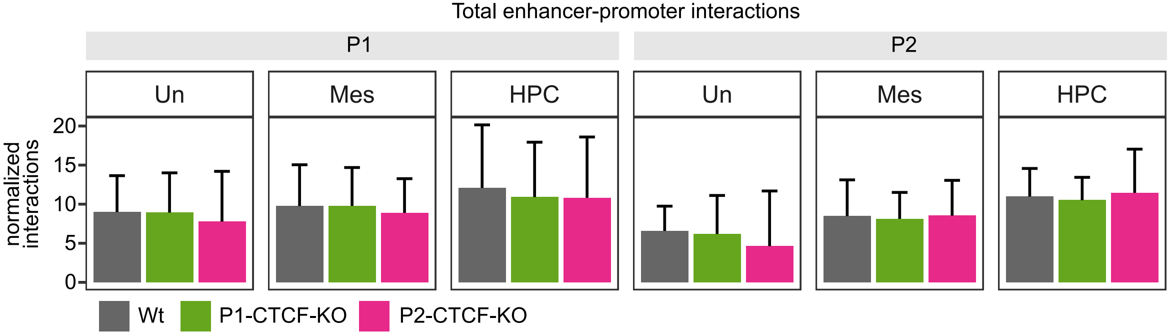
Total enhancer-promoter interactions
P1
P2
Un
Mes
HPC
Un
Mes
HPC
normalized interactions
0
5
10
15
20
Wt
P1-CTCF-KO
P2-CTCF-KO

### Supplementary Figure 12

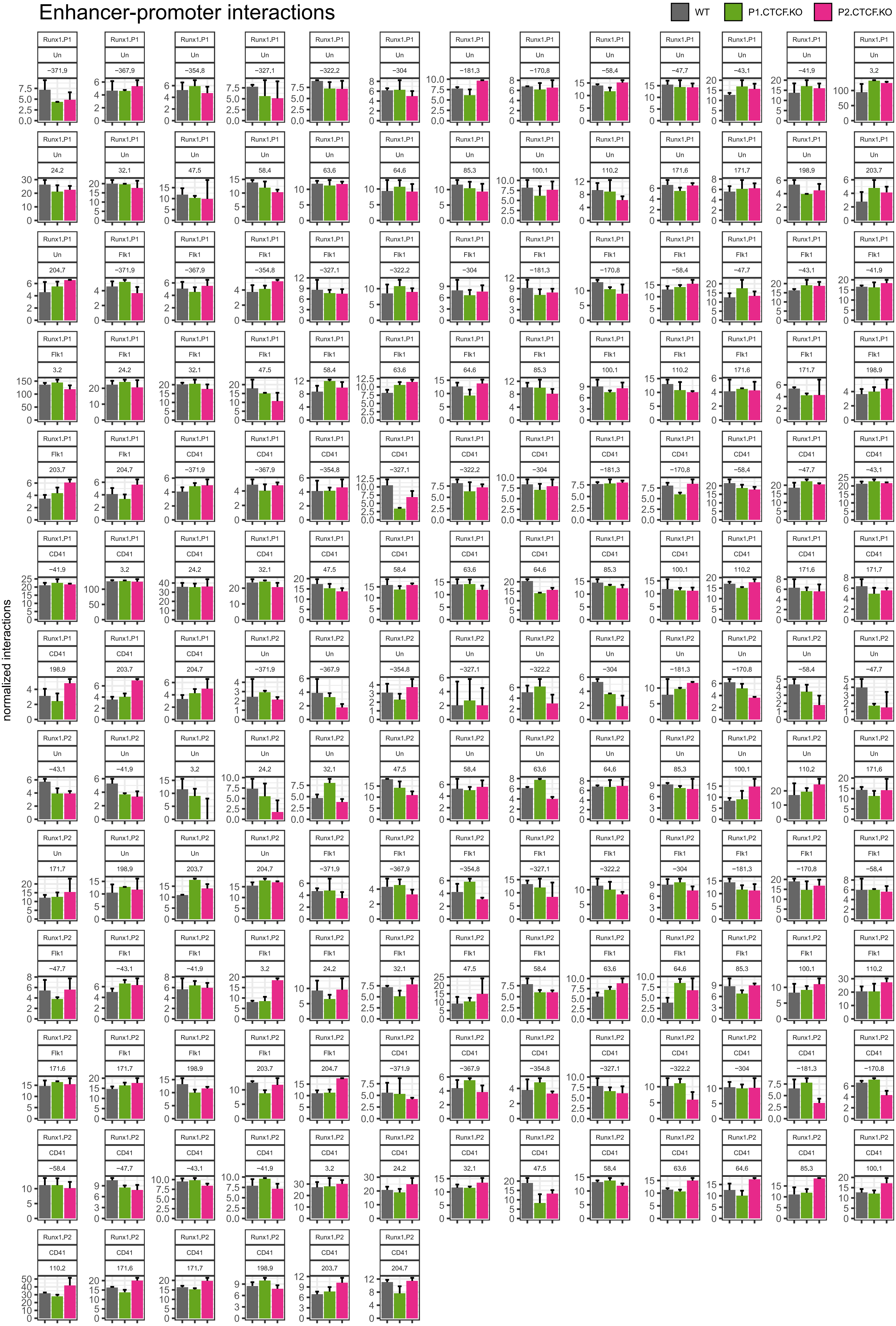

**Supplementary Table 1 - List of open chromatin sites in the *Runx1* TAD**

| mm9 | start | stop | Undiff | Meso | HPC | distance to Runx1 ATG | Known element overlap | ref |
|---|---|---|---|---|---|---|---|---|
| chr16 | 92498792 | 92498856 | | Y | | 327.1 | | |
| chr16 | 92498976 | 92499057 | Y | | | 326.9 | | |
| chr16 | 92515194 | 92515332 | Y | | | 310.6 | | |
| chr16 | 92520183 | 92520297 | Y | Y | | 305.6 | | |
| chr16 | 92544652 | 92544733 | Y | | | 281.2 | | |
| chr16 | 92566069 | 92566274 | Y | Y | | 259.7 | | |
| chr16 | 92566562 | 92566632 | | Y | | 259.3 | | |
| chr16 | 92571567 | 92571768 | Y | | | 254.2 | | |
| chr16 | 92605770 | 92605858 | Y | | | 220.1 | | |
| chr16 | 92606055 | 92606271 | Y | | | 219.7 | | |
| chr16 | 92621095 | 92621162 | | | Y | 204.8 | +204 | Schutte et al. 2016 |
| chr16 | 92632338 | 92632592 | | Y | | 193.4 | | |
| chr16 | 92639468 | 92639526 | Y | | | 186.4 | | |
| chr16 | 92652205 | 92652314 | | Y | | 173.6 | | |
| chr16 | 92680959 | 92681017 | Y | | | 144.9 | | |
| chr16 | 92686184 | 92686312 | Y | | | 139.6 | | |
| chr16 | 92697559 | 92697773 | Y | Y | Y | 128.2 | P2 | Levanon and Groner, 2004 |
| chr16 | 92698058 | 92698142 | | Y | | 127.8 | P2 | Levanon and Groner, 2004 |
| chr16 | 92698987 | 92699142 | Y | Y | | 126.8 | | |
| chr16 | 92709049 | 92709132 | Y | | | 116.8 | | |
| chr16 | 92715843 | 92716073 | | Y | Y | 109.9 | +110 | Schutte et al. 2016 |
| chr16 | 92753499 | 92753649 | | Y | | 72.3 | | |
| chr16 | 92778300 | 92778757 | Y | Y | | 47.4 | | |
| chr16 | 92778949 | 92779007 | Y | | | 46.9 | | |
| chr16 | 92785249 | 92785421 | | Y | | 40.6 | | |
| chr16 | 92787368 | 92787515 | | | Y | 38.4 | | |
| chr16 | 92801791 | 92802205 | | Y | Y | 23.9 | +23 | Nottingham et al. 2007 |
| chr16 | 92819293 | 92819368 | | | Y | 6.6 | | |
| chr16 | 92822529 | 92822841 | | Y | Y | 3.2 | +3 | Schutte et al. 2016 |
| chr16 | 92828323 | 92828494 | | Y | Y | -2.5 | | |
| chr16 | 92828609 | 92828698 | | Y | | -2.8 | | |
| chr16 | 92857265 | 92857329 | | Y | | -31.4 | | |
| chr16 | 92867588 | 92867655 | | | Y | -41.7 | -42 | Schutte et al. 2016 |
| chr16 | 92874454 | 92874521 | | | Y | -48.6 | | |
| chr16 | 92884073 | 92884489 | | Y | Y | -58.4 | -59 | Schutte et al. 2016 |
| chr16 | 92892091 | 92892223 | | Y | | -66.3 | | |
| chr16 | 92996475 | 92996601 | | Y | | -170.6 | -171 | Harland et al. 2021 |
| chr16 | 92997104 | 92997446 | | Y | | -171.4 | -171 | Harland et al. 2021 |
| chr16 | 92999470 | 92999537 | Y | | | -173.6 | | |
| chr16 | 93006972 | 93007130 | Y | Y | | -181.2 | -181 | Harland et al. 2021 |
| chr16 | 93020651 | 93020717 | Y | | | -194.8 | | |
| chr16 | 93041725 | 93041882 | Y | | | -215.9 | | |
| chr16 | 93047887 | 93048252 | Y | | | -222.2 | | |
| chr16 | 93071978 | 93072322 | | Y | | -246.3 | | |

| | | | | | | | | |
|---|---|---|---|---|---|---|---|---|
| chr16 | 93103389 | 93103447 | Y | | | -277.5 | | |
| chr16 | 93129851 | 93130048 | | Y | | -304.1 | -303 | Marsman et al. 2017 |
| chr16 | 93137686 | 93137744 | Y | | | -311.8 | | |
| chr16 | 93137785 | 93137891 | | Y | | -311.9 | | |
| chr16 | 93147823 | 93148052 | | Y | | -322 | -321/322 | Schutte et al. 2016 |
| chr16 | 93153415 | 93153690 | | Y | | -327.7 | -327/328 | Schutte et al. 2016 |
| chr16 | 93159260 | 93159320 | Y | | | -333.4 | | |
| chr16 | 93217970 | 93218034 | | Y | | -392.1 | | |
| chr16 | 93293307 | 93293539 | Y | Y | Y | -467.5 | | |
| chr16 | 93293634 | 93293701 | Y | | | -467.8 | | |
| chr16 | 93329022 | 93329142 | | Y | | -503.2 | | |
| chr16 | 93335261 | 93335343 | | Y | | -509.4 | | |
| chr16 | 93336087 | 93336202 | | Y | | -510.3 | | |
| chr16 | 93346464 | 93346528 | | Y | | -520.6 | | |
| chr16 | 93366653 | 93366773 | | Y | | -540.8 | | |
| chr16 | 93399533 | 93399597 | | Y | | -573.7 | | |
| chr16 | 93438025 | 93438095 | | Y | | -612.2 | | |
| chr16 | 93438201 | 93438272 | | Y | | -612.3 | | |
| chr16 | 93438470 | 93438613 | Y | | | -612.7 | | |
| chr16 | 93444163 | 93444230 | Y | | | -618.3 | | |
| chr16 | 93454640 | 93454730 | Y | | | -628.8 | | |
| chr16 | 93459127 | 93459196 | Y | | | -633.3 | | |
| chr16 | 93459610 | 93459681 | Y | | | -633.8 | | |
| chr16 | 93463746 | 93463906 | Y | | | -637.9 | | |
| chr16 | 93491758 | 93491841 | Y | | | -665.9 | | |
| chr16 | 93511947 | 93512012 | Y | | | -686.1 | | |
| chr16 | 93512384 | 93512457 | Y | | | -686.5 | | |
| chr16 | 93540122 | 93540213 | | Y | | -714.3 | | |
| chr16 | 93544040 | 93544234 | Y | Y | | -718.2 | | |
| chr16 | 93549667 | 93549763 | Y | | | -723.8 | | |
| chr16 | 93549805 | 93549946 | Y | | | -724 | | |
| chr16 | 93550018 | 93550202 | Y | | | -724.2 | | |
| chr16 | 93576908 | 93576984 | Y | | | -751.1 | | |
| chr16 | 93577055 | 93577199 | Y | | | -751.2 | | |
| chr16 | 93586814 | 93587022 | Y | | | -761 | | |
| chr16 | 93588107 | 93588171 | | Y | | -762.2 | | |
| chr16 | 93591267 | 93591381 | Y | | | -765.4 | | |
| chr16 | 93598152 | 93598255 | Y | Y | Y | -772.3 | | |
| chr16 | 93603965 | 93604398 | Y | Y | Y | -778.3 | | |
| chr16 | 93607994 | 93608092 | Y | | | -782.2 | | |
| chr16 | 93615876 | 93616012 | Y | Y | | -790.1 | | |

**Supplementary Table 2 - List of previously published *Runx1* enhancers**

| Distance to *Runx1* ATG (in kb) | mm9 coordinates *chr* | *start* | *stop* | Reference |
|---|---|---|---|---|
| -371 | chr16 | 93197216 | 93198301 | Marsman et al. 2017 |
| -368 | chr16 | 93193448 | 93194216 | Marsman et al. 2017 |
| -354 | chr16 | 93180228 | 93181138 | Marsman et al. 2017 |
| -327/328 | chr16 | 93152333 | 93153713 | Schutte et al. 2016; Marsman et al. 2017 |
| -321/322 | chr16 | 93147786 | 93148307 | Schutte et al. 2016; Marsman et al. 2017 |
| -303 | chr16 | 93129687 | 93130187 | Marsman et al. 2017 |
| -181 | chr16 | 93006325 | 93007986 | Harland et al. 2021 |
| -171 | chr16 | 92996191 | 92997200 | Harland et al. 2021 |
| -59 | chr16 | 92883870 | 92884686 | Schutte et al. 2016; Marsman et al. 2017 |
| -48 | chr16 | 92873240 | 92873850 | Marsman et al. 2017 |
| -43 | chr16 | 92868720 | 92869213 | Schutte et al. 2016 |
| -42 | chr16 | 92867258 | 92868232 | Schutte et al. 2016 |
| +3 | chr16 | 92822418 | 92822899 | Schutte et al. 2016 |
| +23 | chr16 | 92801742 | 92802272 | Nottingham et al. 2007; Bee et al. 2010; Ng et al. 2010; Schutte et al. 2016; Marsman et al. 2017 |
| +24 | chr16 | 92801109 | 92801727 | Nottingham et al. 2007; Schutte et al. 2016 |
| +32 | chr16 | 92792657 | 92794853 | Ng et al. 2010 |
| +59 | chr16 | 92767120 | 92767941 | Ng et al. 2010 |
| +64 | chr16 | 92760779 | 92761825 | Ng et al. 2010 |
| +87 | chr16 | 92739637 | 92741585 | Ng et al. 2010 |
| +99 | chr16 | 92725390 | 92726237 | Ng et al. 2010 |
| +110 | chr16 | 92715159 | 92716303 | Schutte et al. 2016; Marsman et al. 2017 |
| +171 | chr16 | 92653987 | 92654487 | Cauchy et al. 2015; Fitch et al. 2019 |
| +199 | chr16 | 92626889 | 92627053 | Ortt et al. 2008 |
| +204 | chr16 | 92620882 | 92621464 | Schutte et al. 2016 |

**Supplementary Table 3 - List of Antibodies**

| *Target* | *Fluorochrome* | *Manufacturer* | *Catalog #* | *Purpose* |
|---|---|---|---|---|
| Flk1 | APC | eBioscience | 17-5821-81 | FACS |
| CD41 | PE | BD Pharmigen | 558040 | FACS |
| CD45 | APC-eflour780 | eBioscience | 13-5821-81 | FACS |
| Ter119 | PE-Cy7 | BD Pharmigen | 557853 | FACS |
| VE-cadherin | APC | eBioscience | 17-1441-82 | FACS |
| CD41 | unconjugated | BD Pharmigen | 553847 | ICC |
| Runx1/2/3 | unconjugated | Abcam | ab92336 | ICC |
| CD31 | unconjugated | R&D | AF3628 | ICC |
| anti-Rat | Alexa Fluor 555 | Invitrogen | A-21434 | ICC |
| anti-Goat | Alexa Fluor 647 | Invitrogen | A-21447 | ICC |
| anti-Rabbit | Alexa Fluor 647 | Invitrogen | A-11008 | ICC |
| CTCF | unconjugated | EMD Millipore | 07-729 | ChIP |

**Supplementary Table 4 - List of single guide RNAs**

| *Target site* | *5'-3' sequence* |
|---|---|
| P2-CTCF sgRNA1 | GACTGATCCTCGCGCCGTCG |
| P2-CTCF sgRNA2 | AGCCCCGACATGACCGTGAA |
| P1-CTCF sgRNA1 | GGGTCTCCATAGGGCAAGGC |
| P1-CTCF sgRNA2 | GAGTCCTGTGATGATAGTCA |

#### Supplementary Table 5 - List of primers

| *Target site* | *5'-3' sequence* | *Orientation* | *Purpose* |
|---|---|---|---|
| P2-CTCF | CCTTACTTCTCTTGGGCCTTG | F | 500 bp genotyping PCR |
| P2-CTCF | CTGGTGGCCACTTCCTAATG | R | 500 bp genotyping PCR |
| P2-CTCF | GAAGTGGCACCGAGTCATTTA | F | 1.8 kb genotyping PCR |
| P2-CTCF | CCTGATCGAGCTTCGAACTAAC | R | 1.8 kb genotyping PCR |
| P2-CTCF | CTCGTTTGCATAGAGGAGAC | F | 3 kb genotyping PCR |
| P2-CTCF | CAGTTAGCCAGTCACGTAAG | R | 3 kb genotyping PCR |
| P2-CTCF | GCTACTAATGTATGTGCTCGT | F | 5 kb genotyping PCR |
| P2-CTCF | GCTCATGGTGTGTTAGAGTC | R | 5 kb genotyping PCR |
| P2-CTCF | CGAAACAGGAATCGAGAGAC | F | ddPCR 5'1kb |
| P2-CTCF | GGCAGCTTTGTGTCCAG | R | ddPCR 5'1kb |
| P2-CTCF | CCCTCCACTTTCATCTGTG | F | ddPCR 5'100bp |
| P2-CTCF | CCACAGCTTCTTCCTCTTC | R | ddPCR 5'100bp |
| P2-CTCF | GTTAGGCGTCCTGGGAA | F | ddPCR LOA |
| P2-CTCF | CACAAGATCGACCCTAAGGA | R | ddPCR LOA |
| P2-CTCF | GGGACAGACATTAGGAAGTG | F | ddPCR 3'100bp |
| P2-CTCF | CTACCACCGGTCTGAGAG | R | ddPCR 3'100bp |
| P2-CTCF | CACTTGACACGCACTTGAAA | F | ddPCR 3'1kb |
| P2-CTCF | CCCTCGGTAGAGTCCCA | R | ddPCR 3'1kb |
| P1-CTCF | ACTTAAGTGTCCACTCCGATTA | F | 500 bp genotyping PCR |
| P1-CTCF | GGGATTAAGCACTTCTTTAGGC | R | 500 bp genotyping PCR |
| P1-CTCF | CCACCTATTGACCTCTTCGTTC | F | 1.8 kb genotyping PCR |
| P1-CTCF | TGCTACTGACTAATTTGAGGGTATT | R | 1.8 kb genotyping PCR |
| P1-CTCF | CGAGCTCCACTCAAAGAAAT | F | 3 kb genotyping PCR |
| P1-CTCF | TCTAGGAAGGTCATGGAAATAAG | R | 3 kb genotyping PCR |
| P1-CTCF | TACTCACCTCTCATGAAGCA | F | 5 kb genotyping PCR |
| P1-CTCF | CCTTCTGCACAGAATGTCAA | R | 5 kb genotyping PCR |
| P1-CTCF | GCATGGACATCTCTTGGTAA | F | ddPCR 5'1kb |
| P1-CTCF | CTCCTTTGCTCTCCACAAA | R | ddPCR 5'1kb |
| P1-CTCF | CCTGTAGTCATTTCACTTCAGAAA | F | ddPCR 5'100bp |
| P1-CTCF | GGGCATGAGGACGCTTA | R | ddPCR 5'100bp |
| P1-CTCF | GTCACCTCTGGGTCCTG | F | ddPCR LOA |
| P1-CTCF | GGACTTCCATGACTATCATCAC | R | ddPCR LOA |
| P1-CTCF | CCAAGGAAGCAGCAGTTAAA | F | ddPCR 3'100bp |
| P1-CTCF | CAGCCACTGAATCTCTCCTA | R | ddPCR 3'100bp |
| P1-CTCF | GCTTTGTGGACTGAACAGA | F | ddPCR 3'1kb |
| P1-CTCF | CGACTCTAAGTGTCCAGAGA | R | ddPCR 3'1kb |
| chr4_internal_control | GCAGGCTTGAGTAAGAAGAAG | F | ddPCR internal control |
| chr4_internal_control | ATTTCCCTCTGAGTCTCCTG | R | ddPCR internal control |
| chr1_diploid_control | GGTAGTATCTCCACCGATGA | F | ddPCR diploid control |
| chr1_diploid_control | CTATACTGAAGCGCATGGAC | R | ddPCR diploid control |
| chr6_diploid_control | AGTAGCTAGCCGTTGGTAATAG | F | ddPCR diploid control |
| chr6_diploid_control | GCACTGTGAGGGAAACAAAC | R | ddPCR diploid control |
| chr7_diploid_control | TATAGGTGCAACCCGGAAT | F | ddPCR diploid control |
| chr7_diploid_control | TCGGCCGACTAGTCTTTAG | R | ddPCR diploid control |
| CTCF_positive_control_1 | GGCCAAGATAGAGATGGGTTG | F | ChIP qPCR |
| CTCF_positive_control_1 | GTGCCTGATGCCACCTATAC | R | ChIP qPCR |
| CTCF_positive_control_2 | CAATTACCAACTCCGTTCCT | F | ChIP qPCR |
| CTCF_positive_control_2 | ATTGGTAGAACGACTTTCCG | R | ChIP qPCR |

| | | | |
|---|---|---|---|
| ChIP_negative_control_1 | AAGGCTGAAATGCGGATAAA | F | ChIP qPCR |
| ChIP_negative_control_1 | CCACTTTCCAGCTCTAGGTA | R | ChIP qPCR |
| ChIP_negative_control_2 | GGAATGATCACGTCCTTAGC | F | ChIP qPCR |
| ChIP_negative_control_2 | GAGATGTGTTGGGTCTGTAAA | R | ChIP qPCR |
| DpnII_cut_site | GTGTCACCAAAACCAGCTCA | F | 3C library digestion efficiency qPCR |
| DpnII_cut_site | CCTGGAATCCTTTGGCTCAAG | R | 3C library digestion efficiency qPCR |
| DpnII_cut_site | GGGCAGCTAAGATGCAAGTC | Probe | 3C library digestion efficiency qPCR |
| Uncut_genomic_control | TGGAGGGCATATAAGTGCTACTTG | F | 3C library digestion efficiency qPCR |
| Uncut_genomic_control | TGCTTTTGTCTTCCCCAGAGA | R | 3C library digestion efficiency qPCR |
| Uncut_genomic_control | TGCAGGTCCAAGACACTTCT GATTCTGACA | Probe | 3C library digestion efficiency qPCR |
